# Evolutionary transcriptomics reveals longevity mostly driven by polygenic and indirect selection in mammals

**DOI:** 10.1101/2023.01.09.523139

**Authors:** Weiqiang Liu, Pingfen Zhu, Meng Li, Zihao Li, Yang Yu, Gaoming Liu, Juan Du, Xiao Wang, Jing Yang, Ran Tian, Inge Seim, Alaattin Kaya, Mingzhou Li, Ming Li, Vadim N. Gladyshev, Xuming Zhou

## Abstract

The maximum lifespan varies more than 100-fold in mammals. This experiment of nature may uncover of the evolutionary forces and molecular features that define longevity. To understand the relationship between gene expression variation and maximum lifespan, we carried out a comparative transcriptomics analysis of liver, kidney, and brain tissues of 106 mammalian species. We found that expression is largely conserved and very limited genes exhibit common expression patterns with longevity in all the three organs analyzed. However, many pathways, e.g., “Insulin signaling pathway”, and “FoxO signaling pathway”, show accumulated correlations with maximum lifespan across mammals. Analyses of selection features further reveal that methionine restriction related genes whose expressions associated with longevity, are under strong selection in long-lived mammals, suggesting that a common approach could be utilized by natural selection and artificial intervention to control lifespan. These results suggest that natural lifespan regulation via gene expression is likely to be driven through polygenic model and indirect selection.

## Introduction

Over ∼150 million years of evolution, mammals have diversified dramatically (over 100-fold) in terms of maximum lifespan (hereafter ‘lifespan’ and often used as a proxy for longevity). This experiment of nature has attracted much interest from biologists (*1*). The identification of species lifespan-related genetic variation has been a key approach to resolve this question, with focus primarily on exceptionally long-lived species. For example, a survey of the genome of the bowhead whale (the longest-lived mammal known with a lifespan exceeding 200 years) revealed specific sequence changes in genes associated with DNA repair, cell cycle, and aging (*2, 3*). Naked mole rats, which are the longest-lived rodents (lifespan > 30 years), were reported to harbor unique variations in genes related to macromolecular degradation, mitochondrial function, and telomere maintenance, as well as tumor suppression (*4*). Similarly, substitutions in genes related to the GH/IGF-1 axis are found in the Brandt’s bat, which is the longest-lived flying mammal known (*5*). Recent studies have also shown that elephants could be an attractive model organism to study aging, as they exhibit a long lifespan (> 50 years), a low cancer rate, and present an unexpected expansion of potentially functional TRP53 pseudogenes (*6, 7*). These studies suggest that there is a diversity in genetic factors that supports molecular mechanisms of longevity in mammals.

In addition to genetic variation, the lifespan of mammals is also likely to be modulated by the expression level of genes (*8*). For example, it was shown that IGF1R knockout leads to a lifespan increase of 33% and 15.9% in female and male mice, respectively (*9-11*). Similarly, mTOR inhibition in mice increased the median lifespan of female and male mice by ∼25% (*12-14*). In addition, *SIRT6* overexpression increased the median lifespan of male mice by 14.5% (*15*). Comparing gene expression across species is challenging because variables such as developmental stages and environmental factors can mask or distort genuine expression differences. By assuming that gene expression is primarily shaped by stable selection, comparative transcriptomics analyses have been conducted to investigate gene expression patterns across species from an evolutionary perspective (*16-23*). For example, previous studies compared gene expression in the liver, kidney and brain tissues of 34 mammalian species (*16*) and cultured fibroblast cells of 16 mammals (13 rodents, two bats, and a shrew), revealing a number of genes and pathways showing association with maximum longevity (*17*). These studies found that the expression of genes related to central energy metabolism, DNA damage repair, sugar metabolism, and DNA repair was positively associated with longevity, whereas gene expression associated with mitochondrial metabolism, transcriptional regulation, calcium-mediated signaling pathways, protein ubiquitination, and protein localization was negatively associated (*16, 17*). At the same time, in the comparison of age-related transcriptomic changes in *Myotis*, human, mouse and wolf. *Myotis* exhibits unique molecular mechanisms for lifespan extension in functions related to DNA repair, autophagy, immunity, and tumor suppression (*24*). These studies provide many insights into the relationship between the variation of gene expression and longevity traits across species. Nevertheless, the number of species analyzed in previous studies is relatively small and may not fully represent the diversity of gene expression in mammals.

To better characterize the expression profile of protein-coding genes across the mammalian phylogeny, we generated or obtained transcriptome data from the brain, kidney and liver tissues of 106 mammals, covering 16 orders and 45 families. First, gene expression in different organs and species-specific expression patterns were assessed; then genes whose expression levels was significantly correlated with longevity traits were identified within a phylogenetic framework. Pathways that showed signatures of expression changes associated with longevity were identified using a modified summary approach. Finally, an integrated analysis of gene expression and selection pressure was carried out to measure the intensity of selection of the associated genes. The data and analyses presented in this study currently represent the most comprehensive characterization of gene expression in mammalian organs and contribute to our understanding of how lifespan is regulated at the level of gene expression.

## Results and Discussion

### Data generation and species-specific gene expression

To capture the diversity of gene expression across mammals, RNA-seq data (∼5.2 billion Illumina NovaSeq 6000 reads) were generated from polyadenylated RNA fraction of liver and kidney tissues of 56 species (**Table S1, Fig. 1A, Fig. S1**, and **Methods**). Previously published transcriptomes of liver, kidney and brain of 50 additional species were also used (*2, 16, 18, 25-35*) (**Fig. 1A, Fig. S1, Table S1**, and **Methods**). After filtering and orthologs calling, a comprehensive expression dataset was obtained for 13,508 protein-coding genes in three organs of 106 species, which covered the following orders: Artiodactyla (*n* = 9), Carnivora (*n* = 12), Chiroptera (*n* = 36), Cingulata (*n* = 1), Eulipotyphla (*n* = 5), Hyracoidea (*n* = 1), Lagomorpha (*n* = 1), Perissodactyla (*n* = 1), Pilosa (*n* = 1), Primates (*n* = 16), Rodentia (*n* = 18), and Scandentia (*n* = 1). In addition to 102 placental mammals, our dataset included the platypus (Monotremata), the Tasmanian devil (Dasyuromorphia), an opossum (Didelphimorphia), and the sugar glider (Diprotodontia) (**Fig. 1A** and **Table S2**). Information on adult weight (AW) and longevity-related traits— including maximum lifespan (ML), female time to maturity (FTM), adult-weight-adjusted residuals (i.e., MLres and FTMres), and other life-history traits (habitats, feeding habit, etc.) were also collected and analyzed (see **Methods, Fig. 1A** and **Table S2**). Among these traits, ML and FTM reflect changes in absolute longevity, while the residuals indicate changes in relative longevity (**Fig. 1A**). Three algorithms (i.e., mice, missForest, and Phylopars) were used to impute and estimate missing life-history data for the species analyzed (**Figs. 1B-C, Fig. S2, Table S2**, and **Methods**).

**Fig. 1.**
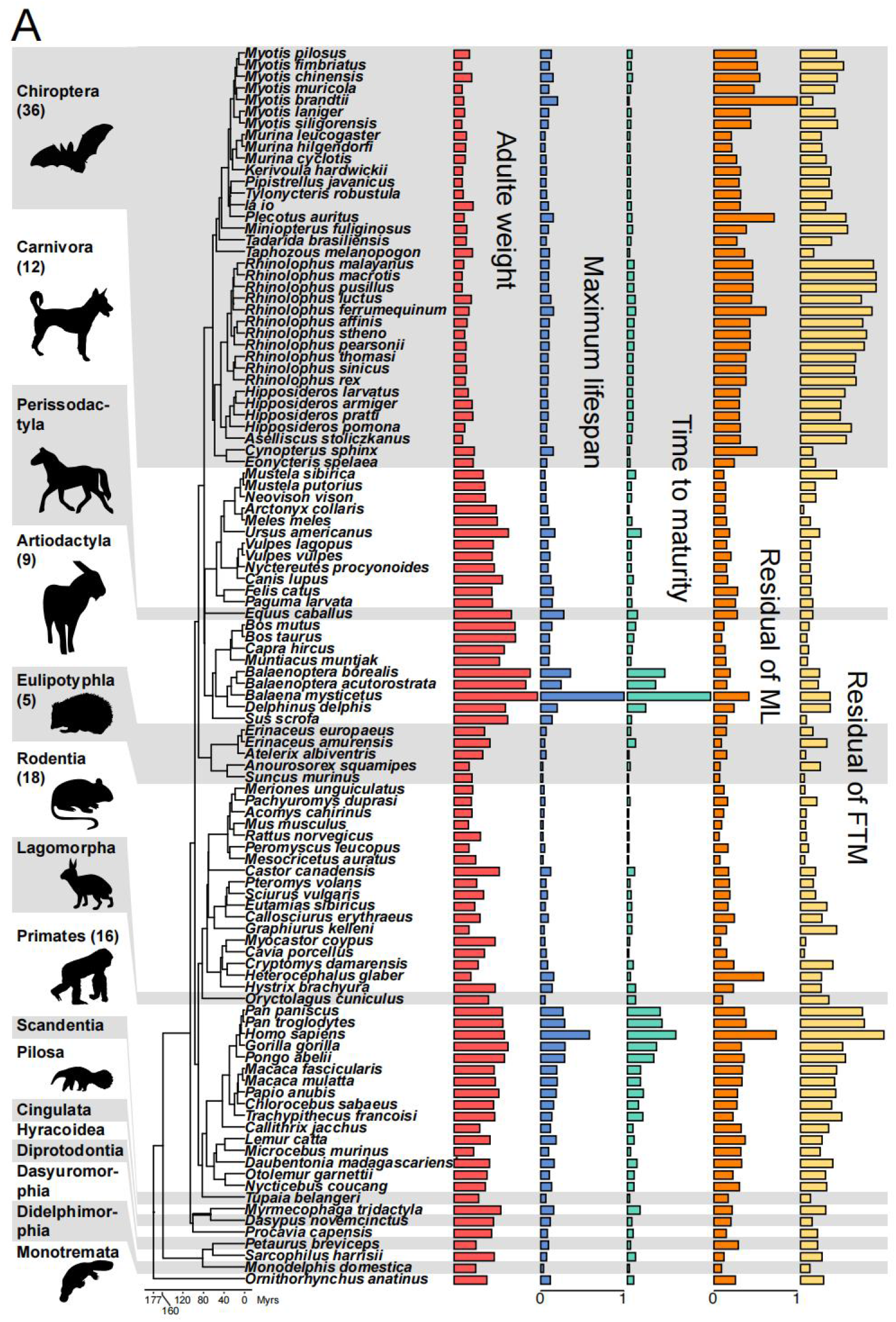
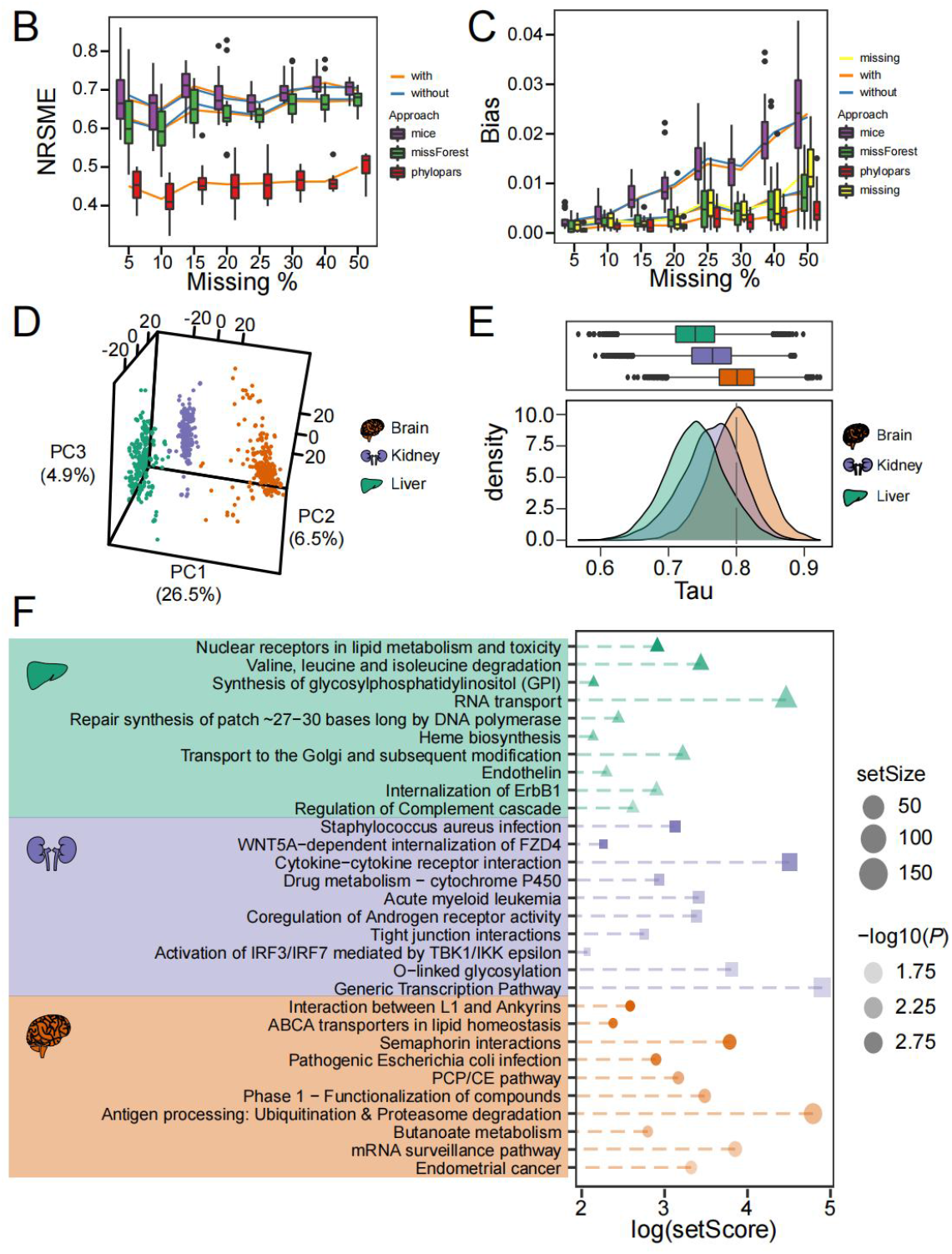
Life-history traits and gene expression profile of mammals. (*A*) Mammalian phylogenetic tree with corresponding life-history traits. From left to right: Adult weight (log_10_-transform), maximum lifespan, female sexual maturity time, and residual of the maximum lifespan and female sexual maturity time relative to the adult weight. Each bar denotes a value of life-history variable for a particular organism in standard scale. The animals image retrieved from PhyloPic (http://www.phylopic.org/) (*B*) Estimation accuracy of missing values of life history traits: the x-axis is the proportion of missing values, and the y-axis is the standard root mean square error (NRSME). (*C*) Estimation bias of missing values: the x-axis is the proportion of missing values, and the y-axis is the bias of biological significance. (*D*) Principal component analysis of gene expression across tissues. The first three principal components (PCs) and their variance explanation percentages are shown. Each repetition is treated as a point. (*E*) Distribution of species-specific expression index (Tau) in the three tissues. Different organs are shown by different colors. The x-axis is the species-specific expression index; the y-axis represents the frequency (below). The dotted line represents the threshold of the species-specific expression index. (*F*) Enrichment analysis of species-specifically expressed genes for each tissue (Liver: green; Kidney: blue; Brain: orange). The x-axis is the log-transformed gene set score. The depth of the color represents the degree of significance of pathway enrichment, and dot size represents the size of the gene set.

Principal component analysis of the expression data metrics was performed to assess the gene expression patterns across species and tissues. Gene expression was tissue-specific rather than lineage-specific (**Fig. 1D**), consistent with previous reports (*16, 18-20*). To characterize genes with species-specific expression patterns, the specificity index (τ; Tau) for gene expression was calculated for each tissue (**Methods**). Tau ranges from 0 to 1 and indicates how broadly (0) or specific (1) a gene is expressed (*36*). None of the 13,508 genes were broadly expressed across species (τ < 0.2) (**Table S3**), which could be due to the large-scale sampling conducted in this study (**Fig. 1E**). Compared to the liver and kidney, the brain presented the highest number of genes showing species-specific expression (6,716, 2,519 and 1,186 genes for the brain, kidney, and liver, respectively) (**Fig. S3**). The gene with the highest species-specific index in the brain was *GRM1* (glutamate metabotropic receptor 1). The primary role of this gene is to protect neurons from apoptosis (*37, 38*). Studies have shown that motor coordination and context-specific associative learning is impeded in mice lacking GRM1 (*39*). It is interesting that *GRM1* is also the most highly expressed gene in bats, because species that specifically express this gene have excellent spatial memory and the ability to recognize individuals (*40*). In the kidney, a detoxification organ, the most highly species-specific genes were *ZNF518A* (zinc finger protein 518A) and *UBE2N* (ubiquitin-conjugating enzyme E2 N). These genes regulate the maintenance of cell types (*41*) and DNA repair (*42, 43*), respectively, suggesting that they play a role in kidney function. *UBE2N* is also highly expressed among long-lived species—such as vervets, naked mole-rats, and greater short-nosed fruit bats—which is consistent with previous reports that DNA repair-related genes are commonly highly expressed in long-lived species (*3, 17, 44*). In the liver, the most species-specific gene was *PTPRG* (protein-tyrosine phosphatase gamma), a marker of oxidative stress associated with inflammation and aging. It has been reported that inflammation caused by obesity can promote *PTPRG* expression in the liver, and an excessive expression of this gene can cause severe liver and insulin resistance (*45*).

To characterize pathways showing expression specificity across organs and species, pathway enrichment analysis was performed under a polygenic model, in which we used the sum of the Tau index of genes in a pathway (*46, 47*) (**Fig. 1F, Tables S4**-**5**, and **Methods**). The brain showed enrichment for “ABCA transporters in lipid homeostasis” (Score = 10.79, *P* = 2.68 × 10^−3^), “semaphorin interactions” (Score = 43.99, *P* = 2.86 × 10^−3^), “PCP/CE pathway” (Score = 23.69, *P* = 8.67 × 10^−3^), and “interaction between L1 and ankyrins” (Score = 13.26, *P* = 1.64 × 10^−3^). Semaphorin interactions and the PCP/CE pathway regulate synaptic formation and help to determine neuronal polarity (*48-51*). They are also important molecular players in the aftermath of nervous system damage events (*52, 53*). In the kidney, specifically expressed genes were enriched in pathways related to detoxification, such as “drug metabolism-cytochrome P450” (Score = 18.80, *P* = 8.03 × 10^−3^), “*Staphylococcus aureus* infection” (Score = 22.77, *P* = 2.24 × 10^−3^), and “cytokine-cytokine receptor interaction” (Score = 90.39, *P* = 3.92 × 10^−3^). This is consistent with the finding above and represents the organ function. In the liver, pathways related to oxidative stress, including “nuclear receptors in lipid metabolism and toxicity” (Score = 18.38, *P* = 1.59 × 10^−3^), “synthesis of glycosylphosphatidylinositol (GPI)” (Score = 8.53, *P* = 3.65 × 10^−3^), and “valine, leucine and isoleucine degradation” (Score = 31.07, *P* = 3.63 × 10^−3^) were significantly enriched. Interestingly, the “valine, leucine and isoleucine degradation” pathway is involved in fatty acid metabolism and immune cell proliferation and is indirectly involved in protein synthesis by affecting the mTOR signaling pathway (*54*). Limiting the intake of these amino acids can extend the lifespan of fruit flies and mice (*55, 56*). These results indicate that mammalian expression profiles are related to organ function.

### Genes with expression level associated with maximum lifespan

Phylogenetic generalized least squares (PGLS) regression analysis was performed to identify genes showing a correlation with longevity between gene expression and four longevity traits (ML, FTM, MLres, and FTMres) and AW (adult weight) within the phylogenetic framework. The ordinary least squares (OLS), Brownian model, and Ornstein-Uhlenbeck model were used for each gene to determine the best correlation (*17, 19, 57*). Based on resampling, the robustness of the correlation was further evaluated through a two-step verification process (see **Methods** for details), to avoid effects by outliers (*P*^*robust*^) or single species (*P*_*max*_) (*17, 19, 58*). Genes that met both a *P*_*robust*_ < 0.01 and *P*_*max*_ < 0.05 threshold were considered significant (**Table 1** and **Table S6**). Based on the obtained results, the total life-history and longevity variation of mammals explained approximately 3.8–4.4% and 2.7– 3.3% of the total variation in gene expression, respectively. This is likely due to the higher number (106) of species analyzed here, compared to a previous report that showed 11%–18% of the inter-species differences explained based on data from 34 mammal species (*16*).

**Table 1.** Statistics on genes whose expression variation is associated with life-history variation.

| Variable* | Liver (n = 13, 508)<sup>†</sup> |  | Kidney (n = 13, 183) |  | Brain (n = 12, 950) |  | Combined<sup>¶</sup> |
| --- | --- | --- | --- | --- | --- | --- | --- |
|  | Nb. of genes<sup>‡</sup> | % from total | Nb. of genes | % from total | Nb. of genes | % from total |  |
| Adult weight | 122 (101) | 0.90 (0.75) | 144 (119) | 1.09 (0.90) | 179 (127) | 1.38 (0.98) | 440 (5) |
| Maximum lifespan | 148 (81) | 1.10 (0.60) | 83 (47) | 0.63 (0.36) | 91 (59) | 0.70 (0.37) | 311 (11) |
| Female time to maturity | 155 (72) | 1.15 (0.53) | 131 (94) | 0.99 (0.71) | 137 (101) | 1.06 (0.69) | 413 (8) |
| Maximum lifespan residual | 96 (51) | 0.71 (0.38) | 96 (56) | 0.73 (0.42) | 131 (81) | 1.01 (0.66) | 323 (1) |
| Female time to maturity residual | 195 (133) | 1.44 (0.98) | 155 (102) | 1.18 (0.77) | 172 (104) | 1.33 (0.80) | 518 (4) |
| Combined<sup>§</sup> | 563 (114) | 4.17 (0.84) | 497 (79) | 3.77 (0.60) | 569 (88) | 4.39 (0.68) | 1557 (275)<sup>#</sup> |

\* The adjusted PGLS P-value cutoff is $P_{robust} < 0.01$ and $P_{max} < 0.05$ .
<sup>†</sup> n denotes total number of orthologous groups assayed in the analysis.
<sup>‡</sup> Number of unique genes associated with trait variation and number of genes specific for a trait (in brackets).
<sup>§</sup> Number of unique genes identified in the organ and number of core genes in the organ (in brackets).
<sup>¶</sup> Number of unique genes identified in three organs for a specific trait and overlap in at least two organs (in brackets).
<sup>#</sup> Number of unique genes identified in three organs for all traits and number of core genes in all organs (in brackets).

We further used the sum of regression coefficient of each gene to identify pathways showing an enrichment with lifespan (*46, 47*) (**Fig. 2A, Fig. S4, Tables S7**, and **Table S8**). Significantly enriched pathways related to the immune system and inflammation were detected in the liver. These included the “inflammatory response pathway” (FTM: Score = 5.54, *P* = 8.00 × 10^−3^; FTMres: Score = 4.36, *P* = 3.02 × 10^−2^), “regulation of IFNG signaling” (ML: Score = 8.62, *P* = 1.90 × 10^−3^; MLres: Score = 7.76, *P* = 2.13 × 10^−3^; FTMres: Score = 4.57, *P* = 2.22 × 10^−2^), “synthesis of leukotrienes (LT) and eoxins (EX)” (MLres: Score = 5.93, *P* = 3.07 × 10^−2^; FTMres: Score = 5.19, *P* = 1.32 × 10^−2^), and “prion diseases” (ML: Score = 10.87, *P* = 7.98 × 10^−3^; FTM: Score = 6.59, *P* = 8.92 × 10^−3^; FTMres: Score = 3.84, *P* = 4.32 × 10^−2^) (**Fig. 2A, Fig. S4** and **Fig. S5**). This enrichment is consistent with previous studies reporting that individuals with longer lifespans show an improved ability to resist inflammation (*59-62*). Interestingly, some well-known aging processes were enriched by genes whose expression showed a significant correlation with longevity. These included “cellular senescence” (ML: Score = −34.10, *P* = 1.52 × 10^−3^; FTM: Score = −23.26, *P* = 8.14 × 10^−3^; MLres: Score = −42.47, *P* = 2.70 × 10^−4^) (**Figs. 2A-B**, and **Fig. S4**), “direct p53 effectors” (ML: Score = 38.00, *P* = 1.05 × 10^−2^; FTM: Score = 24.03, *P* = 1.94 × 10^−2^) (**Figs. 2A-C**, and **Fig. S4**), which also found the same trend in two other tissues. And “Regulation of CDC42 activity” (ML: Score = −11.24, *P* = 3.17 × 10^−3^; FTM: Score = −4.51, *P* = 3.25 × 10^−4^) has the opposite trend in liver and brain (**Fig. 2A, Fig. S4**, and **Fig. S5**). In parallel, pathways related to mitochondrial function were also detected, such as “alpha-linolenic (Ω3) and linoleic (Ω6) acid metabolism” (ML: Score = −6.27, *P* = 6.80 × 10^−3^; FTM: Score = −4.00, *P* = 2.59 × 10^−2^), and “mitochondrial protein import” (ML: Score = −15.53, *P* = 2.95 × 10^−3^; FTM: Score = −10.51, *P* = 1.13 × 10^−2^), which were enriched by genes that showed a significant negative correlation with longevity (**Fig. 2A, Fig. S4**, and **Fig. S5**). In line with this observation, the expression of genes related to central energy metabolism was reported to be downregulated in long-lived animals in a previous study of 34 mammals (*16*). The top genes positively associated with longevity in the liver were *CEP152* (MLres: *P*_*robust*_ = 1.84 × 10^−3^; FTMres: *P*_*robust*_ = 1.91 × 10^−4^), *CACYBP* (ML: *P*_*robust*_ = 3.52 × 10^−4^), *LRP8* (FTM: *P*_*robust*_ = 4.27 × 10^−4^), *RASL12* (FTMres: *P*_*robust*_ = 5.53 × 10^−4^) and *SLC25A12* (ML: *P*_*robust*_ = 2.56 × 10^−4^; FTM: *P*_*robust*_ = 7.21 × 10^−4^) (**Table S6**). Some of them are of interest in relation to aging. For example, the loss of *CEP152* function results in increased DNA damage and delayed entry of the S phase (*63*). *CACYBP* promotes autophagy under starvation conditions (*64*), and counteracts oxidative stress, and maintain nucleolus stability (*65*). *SLC25A12* (encodes aspartate-glutamate carrier isoform 1; AGC1) is highly expressed in liver cancer cells, where it promotes cell division (*66*). Among the top genes negatively associated with longevity in the liver (namely, *NRROS, SCAMP4, RBBP7, MRPL2*, and *F2R*) (**Fig. S8A**), NRROS (FTM: *P*_*robust*_ = 1.35 × 10^−4^) is acetylated after *SIRT6* expression is depleted, and the activity of this gene was found to be correlated with lifespan across mammals (*67*). High expression of *SCAMP4* (FTM: *P*_*robust*_ = 4.52 × 10^−4^) can support senescence-associated secretory phenotype and accelerate aging (*68*). *RBBP7* (ML: *P*_*robust*_ = 1.05 × 10^−3^) was found to be downregulated in senescent cells and is one of the causes of increased DNA damage (*69*). *MRPL2* (ML: *P*_*robust*_ = 1.31 × 10^−3^) activates the mitochondrial unfolded protein response and extends lifespan in worms (*70*). Inhibition of *F2R* (ML: *P*_*robust*_ = 2.14 × 10^−3^) can also delay aging in worms (*71*).

**Fig. 2.**
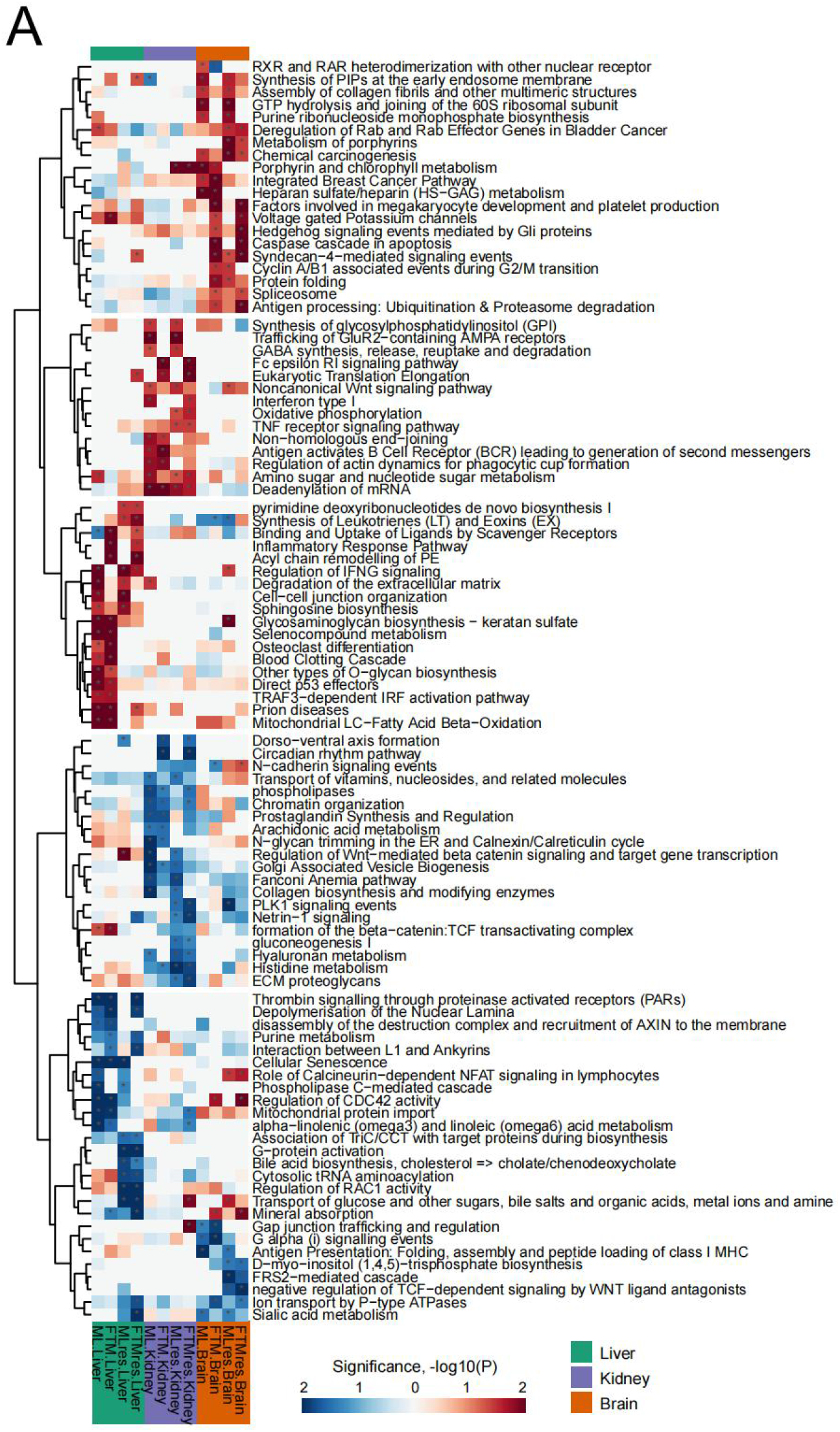
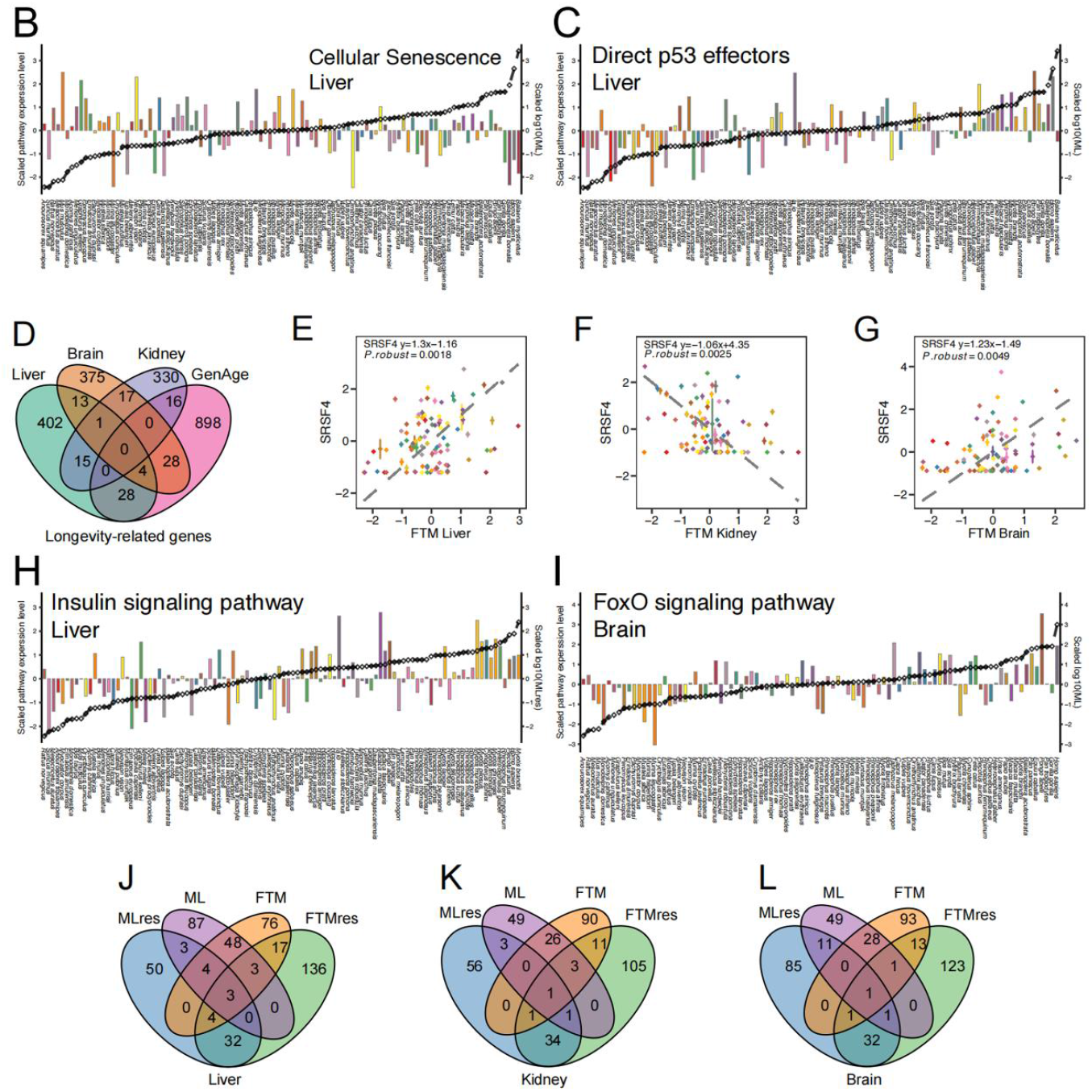
Relationship between gene expression variation and longevity. (*A*) Heat map of the pathways enriched by longevity-related genes. The color intensity indicates the degree of significance, and the *P* value is transformed (-log10). Each row represents a different pathway, and each column represents traits related to longevity (marked at the bottom). Among them, red and blue colors show positive and negative correlations, respectively. The bar graph is the total expression of (*B*) Cellular Senescence negative longevity-related genes and (*C*) Direct p53 effectors positive longevity-related genes in the liver (y-axis on the left). Black line is the relative value of life-history variable (y-axis on the right). Species are shown at the bottom. All values are in standard scale. (*D*) Venn diagrams of the three tissues for longevity-related genes and aging genes of model organisms, obtained from the GenAge database. (*E, F, G*) *SRSF4* is positively correlated with longevity traits in the liver and brain, and negatively correlated in the kidney. In each figure, the y-axis is the scaled expression level of each gene with 0 as the center, and the x-axis represents the longevity traits (ML: maximum lifespan; FTM: female time to maturity; MLres and FTMres: ML and FTM residuals adjusted for adult weight). Potential outliers have been removed. The optimal phylogenetic regression equation, *P* value is included in the figure. The bar graph is the total expression of (*H*) Insulin signaling pathway and (*I*) FoxO signaling pathway positive longevity-related genes in the liver and brain, respectively (y-axis on the left). Black line is the relative value of life-history variable (y-axis on the right). Species are shown at the bottom. All values are in standard scale. (*J, K, L*) Venn diagrams of each tissue for longevity-related genes.

In the kidney, pathways involved in transcription and translation regulation, including “eukaryotic translation elongation” (FTM: Score = 8.26, *P* = 1.67 × 10^−2^; FTMres: Score = 11.09, *P* = 1.44× 10^−3^), and “deadenylation of mRNA” (ML: Score = 8.58, *P* = 9.26 × 10^−3^; FTM: Score = 6.36, *P* = 5.55 × 10^−3^; MLres: Score = 8.95, *P* = 1.70 × 10^−2^; FTMres: Score = 6.19, *P* = 2.46 × 10^−2^), were enriched by genes that were positively correlated with longevity-related traits (**Fig. 2A, Fig S4**, and **Fig. S6**). Interestingly, the “mRNA surveillance pathway” (MLres: Score = −27.32, *P* = 1.60 × 10^−3^) and “tRNA aminoacylation” (FTM: Score = −11.18, *P* = 4.89 × 10^−3^), were enriched by genes negatively correlated with longevity (**Fig. 2A, Fig. S4**, and **Fig. S6**). It has been reported that the regulation of translation fidelity is one of the factors controlling lifespan in a large range of organisms (*72, 73*). For example, naked mole-rat has higher translational fidelity than mouse (*74*). Transfer RNA aminoacylation is inhibited in senescent cells to limit protein synthesis errors (*75*). In addition, seryl-tRNA can directly bind to telomere repeat sequences, leading to telomere shortening and cell senescence (*76*). However, it is unclear why those genes are highly expressed in short-lived species. The “heme biosynthesis” pathway (ML: Score = −4.41, *P* = 3.96 × 10^−2^) was negatively correlated with longevity in the kidney (**Fig. 2A, Fig. S4**, and **Fig. S6**). In contrast, the “porphyrin and chlorophyll metabolism” pathway (MLres: Score = 14.34, *P* = 1.05 × 10^−3^; FTMres: Score = 8.94, *P* = 8.93 × 10^−3^) was positively correlated with longevity in this organ (**Fig. 2A, Fig. S4**, and **Fig. S6**). Heme is an iron porphyrin compound that acts as a precursor for hemoglobin, cytochrome and catalase (*77*). The accumulation of cytochrome promotes cell senescence (*78*), and timely metabolized heme can maintain iron homeostasis and reduce ferroptosis stress in the kidney (*79*). The top genes positively associated with longevity in the kidney included *RHOT2* (ML: *P*_*robust*_ = 3.49 × 10^−4^; FTM: *P*_*robust*_ = 1.72 × 10^−4^), *PARN* (ML: *P*_*robust*_ = 4.00 × 10^−4^), and *DACH1* (MLres: *P*_*robust*_ = 4.25 × 10^−4^); while the negatively associated genes included *BHLHE40* (FTM: *P*_*robust*_ = 1.11 × 10^−4^), *DLAT* (FTMres: *P*_*robust*_ = 4.22 × 10^−4^), and *DIXDC1* (MLres: *P*_*robust*_ = 7.11 × 10^−4^) (**Fig. S8B**). *RHOT2* belongs to the Rho family of GTP enzymes, which are involved in mitochondrial transport and autophagy (*80*), and this gene was positively correlated with longevity in a previous cross-species study (*17*). *DACH1* is involved in suppressing tumor metastasis (*81*), and mutations of this gene were identified in a survey of centenarians (*82*). Among the genes negatively associated with longevity, the overexpression of *BHLHE40* was shown to induce cell senescence, while its knockdown reduced p53-mediated senescence caused by DNA damage (*83, 84*). In addition, BHLHE40 can inhibit fat synthesis mediated by HIF1α under hypoxic conditions (*85*). DLAT can be hydrolyzed by SIRT4, thereby reducing pyruvate dehydrogenase activity and delaying aging (*86, 87*). A study conducted to identify biomarkers of aging, based on RNA-seq and microarray data derived from rats, mice, and humans, showed that the *DLAT* was downregulated during the aging process (*88*).

In the brain, the pathways “metabolism of porphyrins” (MLres: Score = 8.96, *P* = 1.52 × 10^−3^; FTMres: Score = 4.33, *P* = 3.86 × 10^−2^), “glycosaminoglycan biosynthesis–keratan sulfate” (MLres: Score = 7.57, *P* = 2.02 × 10^−3^), “voltage-gated potassium channels” (FTM: Score = 7.34, *P* = 4.01 × 10^−2^; FTMres: Score = 7.33, *P* = 1.31 × 10^−2^), and “syndecan-4 mediated signaling events” (FTM: Score = 5.53, *P* = 1.05 × 10^−2^; MLres: Score = 6.50, *P* = 4.86 × 10^−2^; FTMres: Score = 5.71, *P* =4.11 × 10^−3^) were enriched by genes whose expression positively correlated with longevity traits (**Fig. 2A, Fig. S4**, and **Fig. S7**). Both “metabolism of porphyrins” and “glycosaminoglycan biosynthesis – keratan sulfate,” are associated with clearance of damage in the central nervous system (*89, 90*). Genes in the “voltage-gated potassium channels” pathway have been suggested as potential markers of longevity (*91*), and a deficiency of this pathway results in circadian rhythm disruptions (*92*), shortened lifespan (*93*) and, possibly, obesity (*93, 94*). Among the pathways that were negatively correlated with longevity traits (**Fig. 2A, Fig. S4**, and **Fig. S7**), downregulation of the α2,6-linked sialic acid—which functions within the “sialic acid metabolism” pathway (ML: Score = −8.98, *P* = 3.08 × 10^−2^; MLres: Score = −13.19, *P* = 4.82 × 10^−2^) has been shown to improve cognitive function in mice (*95*), whereas the knockdown of genes in the “PLK1 signaling events” pathway (ML: Score = −15.78, *P* = 9.78 × 10^−3^) induce autophagy, contributing to clearing proteins associated with Alzheimer’s and Parkinson’s diseases (*96*). Among the genes positively related to longevity in the brain, many respond to oxidative stress, including *AMBRA1* (FTMres: *P*_*robust*_ = 4.77 × 10^−4^), *ATG2A* (ML: *P*_*robust*_ = 3.86 × 10^−3^; FTM: *P*_*robust*_ = 4.72 × 10^−4^), and *MCAT* (MLres: *P*_*robust*_ = 9.89 × 10^−4^) (**Fig. S8C**). Both *AMBRA1* and *ATG2A* regulate autophagy and nervous system development (*97*). In addition, *ATG2A* overexpression prolongs the average lifespan in *Drosophila* (*98*). *MCAT* is able to scavenge reactive oxygen species, and its overexpression is neuroprotective (*99*), and reduces age-related oxidative stress in mitochondria (*100*).

### Comparison of longevity-related genes among tissues and gene sets

Unexpectedly, only *SRSF4* (serine/arginine-rich splicing actor 4) showed strong correlation with longevity in all three examined tissues (Liver: ML-*P*_*robust*_ = 4.00 × 10^−4^, FTM-*P*_*robust*_ = 1.80 × 10^−3^; Kidney: ML-*P*_*robust*_ = 7.20 × 10^−2^, FTM-*P*_*robust*_ = 2.50 × 10^−3^; Brain: ML-*P*_*robust*_ = 5.80 × 10^−3^, FTM-*P*_*robust*_ = 4.90 × 10^−3^) (**Figs. 2D-G**). This gene is an essential components of spliceosomes, and thus functions in alternative splicing (*101*), and is downregulated during cellular senescence (*102*). Abnormal SRSF4 function is associated with heart disease and reproductive defects (*103, 104*). Expression changes of splicing factors with age have been described in human and other animal models (*105*). Hence, the strong correlation between longevity and *SRSF4* expression across species indicating that stable alternative splicing could be a ‘long-lived’ feature and that expression level of this gene may be a marker of lifespan. Nevertheless, it is worth noting that the expression of *SRSF4* positively correlated with longevity in the liver and brain, but negatively correlated with longevity in the kidney, suggesting that this gene serves different roles in lifespan control across tissues. It has also been reported that *SRSF4* is negatively regulated by male sex hormones in mice Sertoli cells (*106*) and the kidney is one of the main target organs of these hormones. However, whether the expression pattern of *SRSF4* across tissue is regulated by male hormones, remains unknown to date. Genes (n = 974) with known effects on longevity of model organisms were also examined, revealing that only a few (n = 76, such as *AAK1, KL, NFKB1,STRN*, and *TERT* etc.) showed significant correlation with longevity across phylogeny (*1*) (**Fig. 2D**). This suggests that most of these genes do not serve as a basis for the evolution of longevity across species, although they have been shown to directly contribute to lifespan control in one or more model species.

Nevertheless, several pathways showed correlation with longevity traits in three tissues. The positively related pathways include “Insulin signaling pathway” (**Fig. 2H**), “FoxO signaling pathway” (**Fig. 2I**), “Galactose metabolism”, and “FAS (CD95) signaling pathway”. And the negatively related pathways include “Negative regulation of FGFR signaling pathway”, “Aminoacyl-tRNA biosynthesis”, “Cytosolic tRNA aminoacylation”, “tRNA Aminoacylation”, and “Sema3A PAK-dependent axonal rejection”. Interestingly, “Insulin signaling pathway” (Liver-MLres: Score = 34.45, *P* = 3.46 × 10^−2^; Liver-FTMres: Score = 26.58, *P* = 4.23 × 10^−2^; Kidney-FTM: Score = 23.40, *P* = 3.86 × 10^−2^; Brain-FTM: Score = 27.23, *P* = 4.79 × 10^−2^) was positively correlated with lifespan in all three tissues. We also performed PGLS analyses with longevity traits within Primates, Chiroptera, and Rodentia (**Tables S9**-**S11**). The “Insulin signaling pathway” was positively correlated with longevity in the kidneys of primates (Kidney-ML: Score = 142.62, *P* = 8.03 × 10^−3^; Kidney-FTM: Score = 105.60, *P* = 1.86 × 10^−2^; Kidney-FTMres: Score = 255.45, *P* = 2.76 × 10^−3^), the livers of bats (Liver-ML: Score = 142.62, *P* = 1.38 × 10^−3^; Liver-MLres: Score = 153.41, *P* = 2.58× 10^−4^), and the brains of rodents (Brain-FTM: Score = 69.21, *P* = 3.86 × 10^−2^; Brain-FTMres: Score = 75.28, *P* = 1.74× 10^−2^), but negatively correlated in the kidneys of bats (Kidney-ML: Score = −277.67, *P* = 1.16 × 10^−2^; Kidney-MLres: Score = −160.28, *P* = 4.99× 10^−3^). Though it is reported that inhibiting insulin signaling at the individual level can prolong lifespan (*9-11*), we found that the expression of insulin signaling pathway was higher in animals with longer lifespan, suggesting maintaining of this pathway is critical for regulating aging process. Pathways, such as “Transport of glucose and other sugars, bile salts and organic acids, metal ions and amine compounds” (Liver-MLres: Score = − 29.60, *P* = 1.23 × 10^−2^, Liver-FTMres: Score = −28.74, *P* = 4.00 × 10^−4^; Kidney-FTMres: Score = 21.43, *P* = 1.48 × 10^−2^; Brain-MLres: Score = 23.32, *P* = 3.26 × 10^−2^; Brain-FTMres: Score = 17.72, *P* = 3.65 × 10^−2^) and “Hemostasis” (Liver-MLres: Score = −107.57, *P* = 4.05 × 10^−2^; Kidney-FTM: Score = 75.38, *P* = 2.30 × 10^−2^; Brain-FTM: Score = 90.11, *P* = 2.03× 10^−2^) positively correlated with longevity traits in the kidney and brain, but shown negative correlation in the liver. “G alpha (s) signalling events” (Liver-ML: Score = −21.35, *P* = 1.42 × 10^−2^, Kidney-ML: Score = 21.42, *P* = 3.05 × 10^−2^; Kidney-FTM: Score = 15.37, *P* = 3.70 × 10^−2^; Brain-FTM: Score = 13.43, *P* = 1.74 × 10^−2^; Brain-FTMres: Score = 21.70, *P* = 6.40 × 10^−3^). This is interesting as neither of them showing consistent direction of correlation with longevity traits in three organs. The possible explanation could be the role of pathway in organ aging are different or even opposite. For example, calorie restriction in the kidneys reduces damage from oxidative stress by reducing the activity of certain heme proteins (e.g., Myeloperoxidase, MPO) through anti-inflammatory effects (*107*). However, caloric restriction activates FOXO1 in the liver, which further disrupts mitochondrial function and lipid metabolism via heme (*108*).

To understand the variation of longevity-related genes derived from different genomic sources, we next considered longevity-related genes within metrics that assess the degree of mutation tolerance (essential vs. non-essential genes, and disease-harboring vs. non-disease genes”) and reflect fitness (haplo-sufficient vs. haplo-insufficient genes, and new vs. old genes) (see **Methods** for details). The results (**Table S12**) showed that the direction of the coefficients of absolute (ML and FTM) and relative (MLres and FTMres) longevity-related genes was inconsistent in each gene metric, indicating that most correlations were confounded by body weight. For example, among the mutation tolerance metrics, it was observed that essential genes showed a positive correlation in gene expression with the gradient of ML compared to non-essential genes (average coefficients for the essential/non-essential group: liver, 1.25/0.68; kidney, 0.36/0.17; brain, 0.92/0.31) (**Fig. 3**). However, most essential genes whose expression significantly correlated with MLres showed the opposite trend to those that correlated with ML (average coefficients for essential/non-essential group: liver, −0.31/0.02; kidney, −0.43/0.27; brain, −0.60/−0.91) (**Fig. 3**). The coefficients of young genes and disease-harboring genes had a similar pattern compared to their counterparts. In particular, the coefficient direction of the haplo-sufficient gene in the kidney was always opposite to the other two organs. This observation suggests that more attention should be paid to the selection of organs in aging experiments. The results also showed a considerable variation in the coefficient of haplo-sufficient genes among tissues (**Fig. 3**), which indicates that these genes play diverse roles in lifespan across organs.

**Fig. 3.**
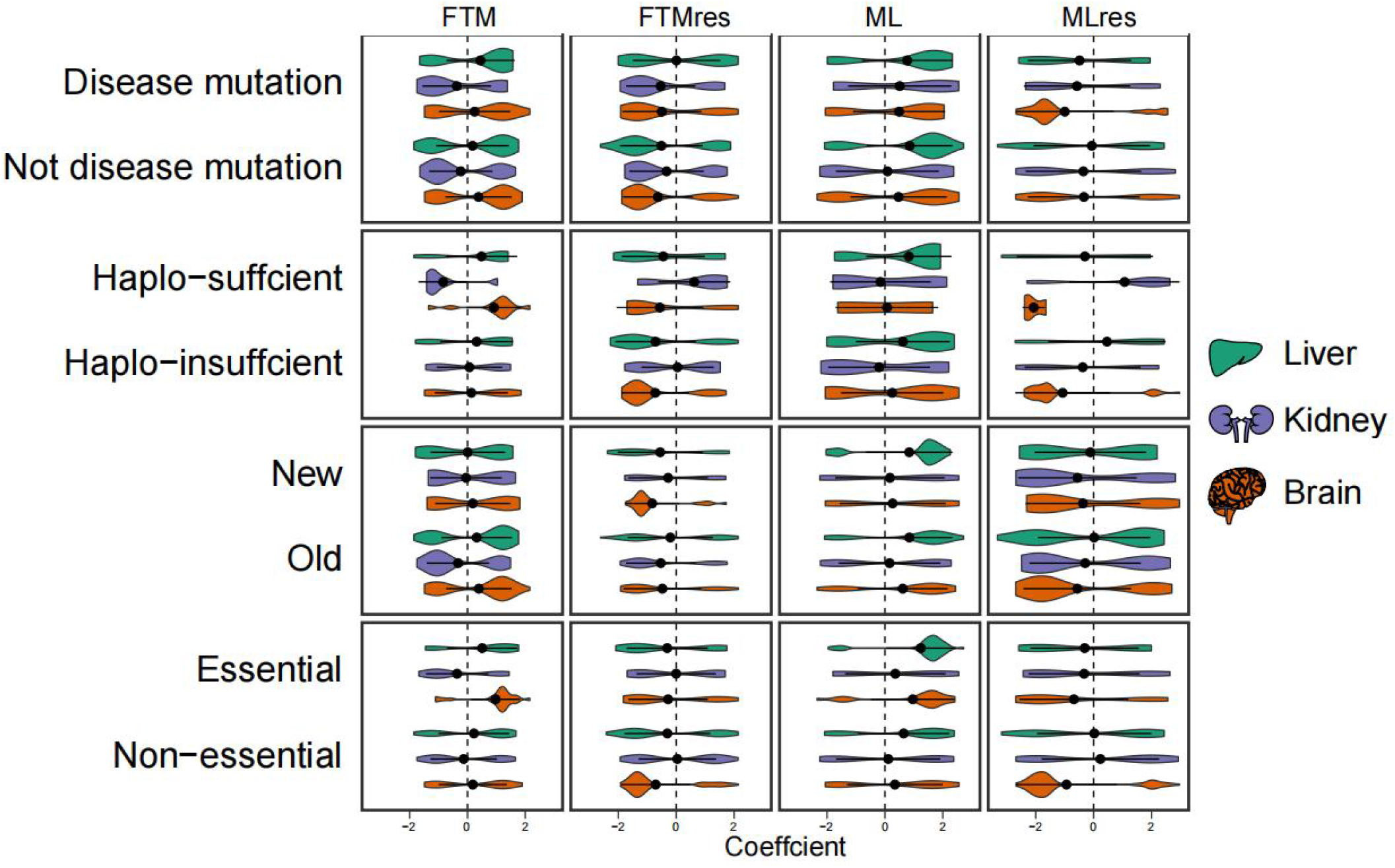
Comparison between the variation of gene expression and the gradient of longevity traits among different gene categories. On the x-axis, the coefficient represents the rate of gene expression variation with life-history gradient. The violin plot and black dots represent the mean of the variation rate in a specific functional gene category, respectively.

### Relationship between selection pressure and gene expression

To uncover the relationship between the intensity of selection and gene expression, we used RELAX in Hyphy (*109*) to assess if selection pressure relaxed by inferring with the parameter (*k*) (**Table S13**). The relaxation parameter *k* was estimated for each gene with long-lived mammals set as the foreground branch (ω_background branch*k*_ = ω_foreground branch_). A *k* > 1 indicates that a gene in the foreground branch is under intensified selection, while *k* < 1 indicates relaxed selection. The genes whose expression was correlated with longevity were divided into four categories (**Fig. 4A**): positively correlated genes under intensified selection (IU), positively correlated genes under relaxed selection (RU), negatively correlated genes under intensified selection (ID), and negatively correlated genes under relaxed selection (RD). The results showed that approximately 62% of total genes are under intensified selection in long-lived mammals (defined as having an ML > 30 yrs.) (**Figs. 4B-C** and **Table S14**). This is interesting, as a previous study reported that genes tend to be under relaxed selection in shorter-lived killifish (*110*). We also tested what type of selection was present at different maximum-lifespan intervals (ML < 12; 12 <= ML <= 26; 26 < ML; 50 < ML), and the results consistently showed that species with longer lifespans tend to have a greater number of genes under intensive selection (**Fig. S9** and **Tables S15**-**17**). We then found that the pattern of selection intensity related to the direction of longevity-associated genes is different across organs (**Figs. 4D-I** and **Fig. S10**). Particularly, in the kidney, a similar proportion of both positively and negatively associated genes was under intensified selection. However, a comparatively higher proportion of positively associated genes—rather than negatively associated genes—was found to be under intensified selection in the liver, while a higher proportion of negatively associated genes was under intensified selection in the brain. It was also observed that several longevity-related genes are under strong intensification of selection in long-lived mammals (**Fig. 4** and **Fig. S10**). For example, among the genes in the IU category, *ICMT* is upregulated in *Drosophila* with extended lifespan (*111*), and *KCNC4*, which mediates the voltage-dependent potassium ion permeability of excitable membranes, is a marker of longevity (*91, 93*). Within the ID category, the downregulation of *CCND2* promotes cell cycle arrest and apoptosis and leads to cell senescence (*112-114*), while *ZNRF2* is one of the top genes negatively correlated with longevity in the liver tissue and is involved in the mTOR signaling pathway (*115*). In addition, we observed that the longevity correlated genes are characterized by either intensified or relaxed positive selection. In parallel, the strength of purifying selection of those genes has been less changed (**Table S14**).

**Fig. 4.**
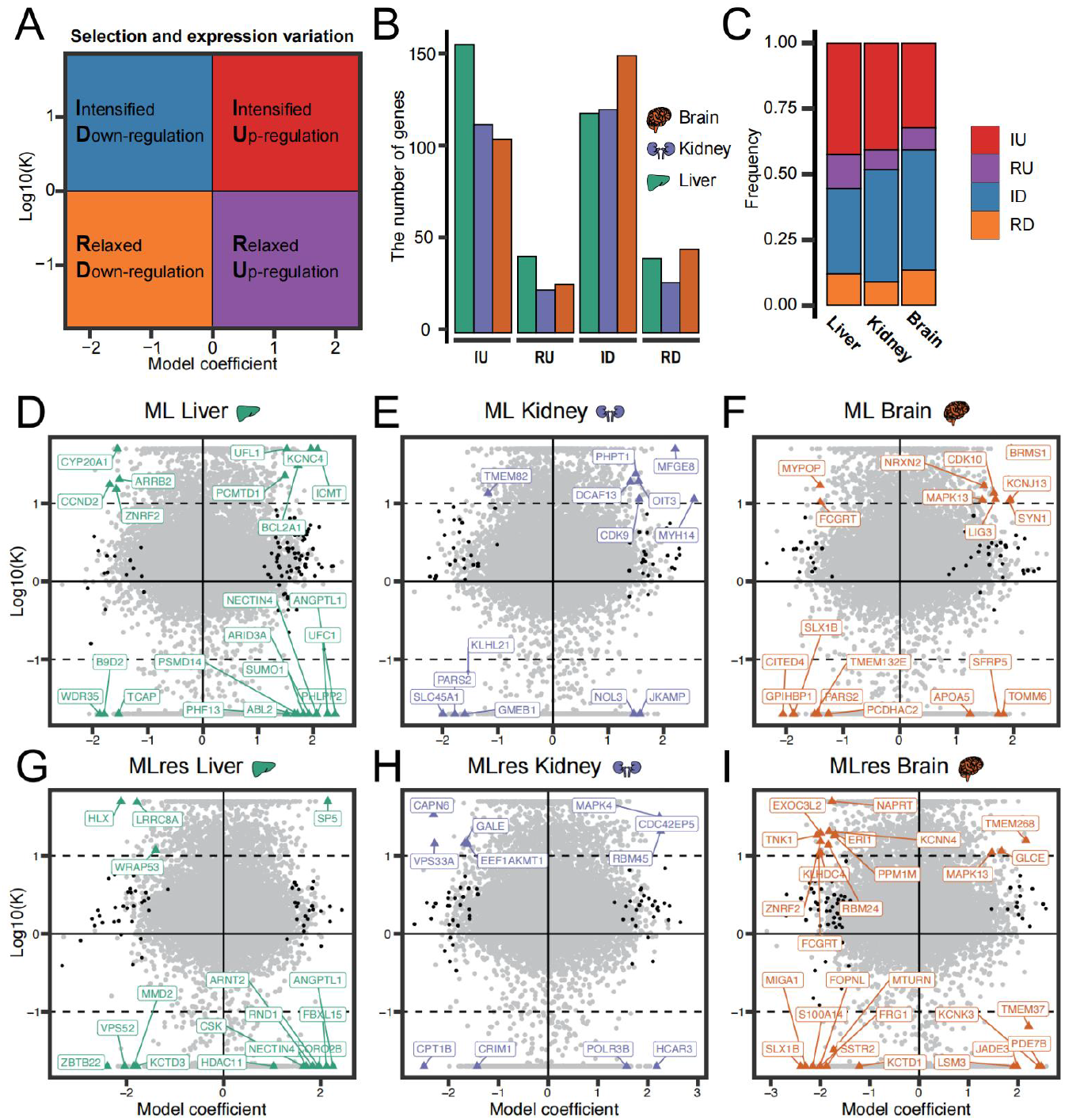
Pattern of association between selection index (*k*) and coefficients, and longevity. (*A*) Colored picture illustrating the annotation of gene classification. (*B*) Number of genes in the three organs assigned to different gene classifications. IU stands for genes that are positively related to longevity and are under intensified selection; RU stands for genes that are positively related to longevity and are under relaxed choice; ID represents a gene that is negatively related to longevity and is under intensified selection; RD stands for genes that are negatively related to longevity and are under relaxed selection. (*C*) Cumulative frequency histogram showing the distribution of gene types in different organs. (*D-I*) Scatter plot showing the log10-transformed relaxation parameter (*k*) on the y-axis, and the variation rate of gene expression along the longevity trait gradient on the x-axis. The colored points represent longevity-related genes with strong selection signals (the color corresponds to the tissue type shown in panel A), the black points represent longevity-related genes with weak selection signals, and the grey points represent genes that are not significant.

Pathway enrichment analysis of longevity correlated genes was performed using the relaxation parameter (*k*) as a statistic to further characterize the cumulative effect of relaxed and intensified selection in long-lived mammals (**Fig. S11** and **Table S18-19**). Pathways related to methionine metabolism were detected, such as the “methionine salvage pathway” (Score = 8.41, *P* = 7.27×10^−4^) and “glycerolipid metabolism” (Score = 21.31, *P* = 2.82×10^−3^), which are enriched by genes under intensified selection. Methionine is one of the essential amino acids, and similarly to calorie restriction, methionine restriction has been reported to extend lifespan (*116-118*), reverse inflammation, and reduce DNA damage (*118, 119*). Interestingly, our analysis revealed that several pathways associated with acceleration of the aging process are under relaxed selection in long-lived mammals. These pathways included “CD28 dependent PI3K/Akt signaling” (Score = −8.31, *P* = 8.69×10^−3^), “WNT ligand biogenesis and trafficking” (Score = −10.42, *P* = 9.69×10^−3^), and the “VEGF signaling pathway” (Score = −7.24, *P* = 4.43×10^−2^). Inhibition of these pathways has been shown to extend lifespan (*120-123*).

## Conclusions

In this study, a comparative transcriptomics analysis of 106 species representative of diverse families was performed to describe gene expression diversity in mammals. It was found that gene expression was conserved, with the strongest specificity in the brain, compared to the liver and kidney, and that the expression of species-specific genes may reflect adaptive traits (e.g., bats’ requirement for long-term memory). In our study, the effect of adult weight on the robustness of longevity-related genes is very limited, and the intersection of longevity-related genes and adult weight-related genes is very small (**Fig. S12A**), which is consistent with previous study (*16, 17*). In addition to very few known aging-related genes (e.g., *TP53, LRP8, RBBP7*, and *SCAMP4*) that showed correlation with longevity, many new genes whose expression levels significantly correlated with longevity across mammalian phylogeny were identified (e.g., *SRSF4* in all tissues, *SLC26A6, ELFN1* and *NEURL1* in the liver, *SCARA3* in the kidney, and *STX5* in the brain) (**Figs. 2D-G** and **Figs. S12B-G**). Enrichment of many well-known aging-related pathways were also observed. These included “Insulin signaling pathway”, “FoxO signaling pathway”, “inflammatory response pathway,” “cellular senescence,” and “p53 signaling pathway”. Importantly, our study also detected other pathways—such as “eukaryotic translation extension,” “non-classical Wnt signaling pathway,” “mRNA polyadenylate,” “mRNA detection pathway,” and “tRNA aminoxylation”. This supports reports that a stable protein synthesis (proteostasis) is important for lifespan control (*73, 124*). We found that longer-lived animals always had more genes that under intensified selection than shorter-lived animals. Although it is generally believed that there is a certain correlation between selection pressure and gene expression (*110*), this was not significantly observed in relation to the gradient of longevity, suggesting lifespan is not favored directly by natural selection (*125*). We still found that methionine restriction is under Intensified selection in long-live mammals. However, given most of our samples are matured males, the variation of gene expression among different ages and genders have not been quantified in our study. And, because of the high-quality genome assembly are not available for many species that analyzed, it is difficult to disentangle the role of the expression changes of paralogous and multi-copy genes in longevity evolution. Nevertheless, the data and results presented in this study would aid future investigations with inclusion of samples at different ages of both genders. Overall, our study suggests that the evolutionary correlation between gene expression and longevity is organ-specific and characterized by polygenic selection. The longevity-associated genes identified could serve as candidate targets for further exploration of healthy aging.

## Materials and Methods

### Tissue collection

The 331 organ samples analyzed in the present study were newly obtained from 56 species (**Tables S1** and **S2**). Liver, kidney and brain tissues were mostly sampled from adult and male individuals, if possible, and were freshly frozen in liquid N_2_ and stored at −80°C. To maximize sample compatibility, each major part of each organ was dissected and homogenized. To objectively detect the biological variation of gene expression, three biological replicates were obtained when possible. All the experimental protocols were approved by the Animal Care and Use Committee of the Institute of Zoology, Chinese Academy of Sciences (No. IOZ-IACUC-2021-129).

### Transcriptome library preparation and sequencing

Total RNA was isolated from frozen tissue using TRIzol^®^ Reagent (Invitrogen). To protect RNA as much as possible during homogenization, we first added 0.2 mL TRIzol^®^ Reagent directly to a tube containing 100 mg of frozen tissue and homogenized using a motorized homogenizer. After homogenization, we added another 0.8 mL of TRIzol^®^ to the tube. The resulting lysate was phase separated with 0.2 ml chloroform and total RNA precipitated with 0.5 ml isopropanol. The RNA was washed twice with 1 ml 75% ethanol and resuspended in DEPC treated ddH2O. The resuspended RNA was assessed for quality (260/280 nm absorbance ratio) and integrity (formaldehyde agarose gel electrophoresis). The sequencing libraries were prepared using the NEBNext Ultra RNA Library Prep Kit for Illumina (NEB, USA), and the transcriptome libraries were sequenced on an Illumina NovaSeq 6000 system (Novogene Co. Ltd) with paired-end reads of 150 bp. NGS QC Toolkit v2.3.3 (*126*) was used to remove reads containing adapters and filter low quality reads (<Q20).

### Orthologous gene sets

Genome annotations (GTF) for 39 mammals with sequenced genomes were obtained from Ensembl, release 99. For the minke whale (*Balaenoptera acutorostrata*), Indian muntjac (*Muntiacus muntjac*), great roundleaf bat (*Hipposideros armiger*), Chinese rufous horseshoe bat (*Rhinolophus sinicus*), Brandt’s bat (*Myotis brandtii*), François’s leaf Monkey (*Trachypithecus francoisi*), and white-footed mouse (*Peromyscus leucopus*) we used GTF annotations and genomes downloaded from NCBI database. For the bowhead whale (*Balaena mysticetus*) we used genomes downloaded from ‘The Bowhead Whale Genome Resource’ (*127*) (**Table S2**). The GTF of bowhead whale was generated using augustus v 2.5.5 (*128*). Draft transcriptome of 59 mammals were *de novo* assembled using trinity v2.11.0 (*129*). First, the RNA-seq reads from same species and tissues were assembled together. Because the Trinity assembler filters low-coverage k-mers, we did not perform quality trim of the reads before assembly. Trinity was run on 150 bp paired-end sequences with default parameters k-mer size of 25 (fixed), minimum contig length of 200, maximum paired fragment length of 500, and adjusted butterfly maximum heap space setting to 30G. To remove redundancy, we then used cd-hit (*130, 131*) to process the assembled transcripts from different tissues of the same species, cluster the sequences with 90% similarity, and leave the longest transcript in each cluster. We used augustus to perform gene prediction on the de-redundant transcripts and obtain GTF annotation files. We used gffread in the cufflink package v2.2.1 (*132*) to extract the CDS sequence, filtered out incomplete ORF transcripts and pseudogene transcripts, and extracted the longest transcript of each gene. Given the genome assembly for most of species are scarce or not well-annotated, multi-copy genes and transcripts were not considered in the analyses. To reduce the effects of paralogs on the ortholog identification, we constructed the human reference sequence using BLAST v2.9.0+ (*133*) to remove highly repetitive and highly similar genes, with e-value < 10^−6^ and Identity > 90% as the filtering threshold. In the end, 18,553 unique protein coding genes were obtained as reference sequences. For other mammals, the longest transcript of each gene was extracted and reciprocal BLAST was performed with the protein sequences from human. The filtering threshold was 10^−6^ for e-value and 30% for identity. Two genes that were best aligned with each other were defined as orthologous genes. When a gene exists in fewer species, it indicates that the gene is not highly conserved and cannot be representative of mammals. However, as the number of species increases, the number of orthologous genes that coexist in all species decreases (only 989 genes are present in all species). To balance the number of species and genes, we filtered out genes that exist in less than 70 species. The final dataset of orthologous gene accounted for 13,916 individual groups of sequences. In downstream analysis, each gene is analyzed individually, and only the species in which the gene is present were considered.

### RNA-seq reads mapping and normalization

Because the complete genome and the *de novo* genome are quite different when compared, we used the CDS sequence of orthologous genes as the reference genome, and generate annotation files in GTF format for RNA-seq data mapping. STAR v2.7.1a (*134*) was used to construct an index. Because of the specificity of the orthologous genomes, the parameters ‘--genomeSAindexNbases’ and ‘--genomeChrBinNbits’ were calculated from the sequence size of the homologous gene set and the read length of different samples. And we used the default parameters to align the RNA-seq data with the orthologous genomes. We used featureCounts v2.0.0 (*135*) to count reads, and eliminate multiple-matched reads (**Table S20)**. Generating gene expression profiles for all species based on pairwise orthologous relationships. Finally for 18,553 genes, abnormally low-expressed genes that is, genes whose expression levels were less than 10 in 4 or more samples were filtered before normalization (2,564 genes were removed). And, abnormally high expression genes, that is, genes whose total expression of all samples accounted for 5% of the expression of the entire data set was also removed (1 gene). The function comBat_seq in the R package sva (*136*) was applied to read counts to remove the batch effect, including the two factors most likely to affect the data: different sources of data (Bioproject and sequencing batches of our data) and the deviation caused by the sequencing platform (*137*). Two factors were adjusted separately. And the covariates were tissue and species. Genes with orthologous in more than 70 species were used for downstream analysis to reduce the false positive rate in the analysis (13,827 genes in total). We calculated the library size of each sample as a normalization factor. The R software package edgeR (*138*) was used to normalize the library size and gene length (based on humans) by log_2_(TMM-RPKM + 1). For paired-end data, featureCounts counts fragments, so calculating RPKM for paired-end data is equivalent to FPKM.

### PCA analysis and species specificity of gene expression

We calculated the variance of each gene on the normalized expression matrix, and selected the top 5,000 genes with the largest variance to perform principal component analysis (PCA) using the R package ‘FactoMineR’ (*139*). In order to define gene sets that are widely expressed by species and species-specifically expressed genes. We calculated the mean value of log2(RPKM-TMM + 1) in each organ for each species. We calculated the species-specific expression index Tau, 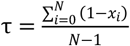, which is used to quantify the tissue specificity of gene expression (*36*). Among them, N is the number of species, *x*_*i*_ is the expression level of the i-th species. Tau > 0.8 is defined as a species-specific gene.

### Life-history data collection and imputation

To accurately estimate the species for which life history data were missing in this study. We collected data on highly correlated life-history traits (AW: adult weight, ML: maximum lifespan, and FTM: female mature time) for a total of 1,250 species from the online databases AnAge (*1*), Animal Diversity Web (https://animaldiversity.org/) and PanTHERIA (*140*), and from the literature. And the phylogenetic tree was retrieved from TimeTree (http://www.timetree.org/) (*141*). Three life-history traits from 816 species were complete and used as a training set and, we employed three imputation methods to estimate the missing data: (i) Based on the Markov chain Monte Carlo method, *mice* introduces the random process into the interpolation process, uses other variables as predictors, and specifies a conditional model for each variable (*142*). We used the predictive mean matching (pmm) as the conditional model in the multiple regression model, or used mean matching (mean) instead if the first run did not converge. We selected the case where the predicted regression score was closest to the missing value. (ii) missForest first uses the mean to interpolate a column of data (*143*) and then uses the remaining variables of the data set to fit a random forest model to estimate missing values by applying trained random forest predictions. This process was looped for all variables that need to be interpolated, and the whole process was repeated until the stop criterion was reached. (iii) Phylopars estimates missing values based on restricted maximum likelihood (*144*). This method calculates the covariance matrix based on phylogenetic and phenotypic components (when multiple trait measures are given). It builds a multivariate normal model that combines the best phylogenetic and phenotypic covariance with the tree to calculate the covariance between the observed and missing values. Therefore, the estimated value was determined by phylogenetic distance (correlation between species) and ectopic relationship (correlation between features).

The percentages of missing values in the missing set were 6% (ML) and 30% (FTM). We tested three types of missing data: (i) Completely missing at random (MCAR); (ii) Missing at random based on weight (MAR.AW), and it was divided into two types of species according to the median weight. Because low-weight species may have more missing values; (iii) Missing at random based on the genetic distance between human (MAR.HD), and it was divided into two types of species according to the half of the farthest genetic distance. Because species with a greater genetic distance from humans may receive less attention from scientists, the life-history is also opted to be missed.

We performed chi-square tests on the two types of MARs in the missing set. In addition, missing values in large-weight species accounted for 17.57% of the total missing values in maximum lifespan (ML), and 82.43% in small-weight species. In FTM, the missing values of the large-weight species accounted for 37.66% of the total missing values, and the small-weight species 62.34%. We introduced multiple missing value ratios (5%, 10%, 15%, 20%, 25%, 30%, 40% and 50%) to the training set to simulate the distribution and pattern of missing values. We used the above three methods for 10 interpolations and imputation. In order to account phylogenetic relationships in the imputation process (Phylopars are only applicable to imputations that include phylogeny), R package PVR was used (*145*) to perform principal coordinate analysis (PCoA) on the genetic distance matrix of 816 species to obtain phylogenetic feature vectors. The phylogenetic relationship after dimensionality reduction is used as other predictor variables in the imputation process. At the same time, we obtained the optimal number for interpolation by adding phylogenetic vectors in the interpolation process incrementally. We evaluated the accuracy of the interpolation based on the normalized root mean square error 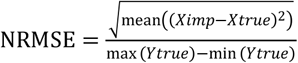.

And, in order to ensure that the estimated value retains biological significance. We also calculated the bias of the slope (Bias = |Slope^*original*^− Slope^*imputed* (*or missing)*^ |) between adult weight and maximum lifespan in the data set after interpolation.

Finally, we selected the best Phylopars based on the evaluation results for the imputation of the complete life history data set (**Table 2**). To identify confounding factors of maximum lifespan, we collected multiple complex effects such as society, diet, habitat, activity, body mass, basal metabolism rate, and offspring per year. We have used MCMCglmm to examine the correlation between longevity and other factors. The result showed that body mass and offspring per year were significantly associated with maximum lifespan, and a weak association between diet and maximum lifespan (**Table S22**). And we also detected a strong correlation between body mass and offspring per year. Previous studies also have shown that the longevity (or female time to maturity) was mainly correlated with body mass (*16*). Nevertheless, many species show with small weight and long maximum lifespan (or female time to maturity). Therefore, we calculated the residuals to correct the confounding effects caused by weight (i.e., MLres and FTMres). Both residual equations are obtained based on linear regression model using the data from the AnAge database (*1*).

### Phylogenetic regression analysis

To identify genes with the expression related to longevity, three evolution models were tested for gene expression in each tissue (mean value of log_2_-scaled TMM-RPKM) and each log_2_-scaled longevity-related trait (ML, FTM, MLres, FTMres), including regression models that do not consider phylogenetic relationships (OLS), and regression models that consider phylogenetic relationships (BM and OU). And the optimal model was selected according to the maximum likelihood methodology. The phylogenetic tree was retrieved from TimeTree (*141*). The unit of branches length of the phylogenetic tree is million years. To avoid randomness, we took a resampling approach (*58*) instead of using conventional P-value corrections (e.g., BH). A two-step method is used to correct the *P* value (*17, 19*). In the first step, the species that has the greatest impact on the slope (i.e., potential outliers) is removed by the residuals (the largest absolute value of the residual is removed), and then the regression is performed again. At this time, the *P* value obtained is defined as *P*_*robust*_ to remove the influence of the outliers on the regression. The second step is to repeat the regression process for the remaining species and remove one of remaining species each time until all remaining species are removed once, and take the largest (least significant) *P* value in the process as *P*_*max*_ to remove the impact of species on regression. The cutoff for identifying longevity-related genes was *P*_*max*_<0.05, *P*_*robust*_<0.01.

To reduce the noise caused by missing data, for each gene, we only consider the species in which the gene exists, and did not add all species to the model, that is, set the expression value of a gene that does not exist in a species to the missing value instead 0.

### Multiple sequence alignment and selection pressure analysis

For each group of orthologous genes, the Perl script ‘translatorX.pl’ (*146*) is used for multiple sequence processing and comparison. This pipeline selects the default parameters of the ‘MAFFT’ (*147*) to first translate the nucleic acid sequence into a protein sequence for multiple sequence alignment and then translate it back into a nucleic acid sequence. We then used ‘GBlock’ (*148, 149*) to select the conservative blocks, with the number of conservative sites in the gene sequence after alignment is at least 75% of the total length of the gene, and the shortest flanking sequence is greater than 85% of the length of the gene after the alignment.

In order to test whether genes are under relaxed selection, we used the minimal model of RELAX (*109*) in ‘Hyphy’, with long-lived animals (ML > 30 years) were set as foreground branches. Because the number of non-long-lived mammals (n=82) is far more than that of long-lived mammals (n=24). We selected 24 representative species with good genome quality from non-long-lived mammals as background branches to eliminate the noise caused by the excessive number of background branch species (**Table S21**). This model uses the likelihood ratio test to compare the two models with the same evolution rate (k = 1) and different rates (k ≠ 1) between the foreground branch and the background branch. The parameters are set to estimate 3 types of ω (ω1: purification selection; ω2: neutral selection; ω3: positive selection). The relaxation parameter k is an index of the selection strength (ω_background branch_ ^k^ = ω_foreground branch_), with k > 1 indicates that the genes in the foreground branch are under intensified selection and k < 1 indicates a relaxed selection. And we also test relaxed selection at different interval (ML < 12; 12 <= ML <= 26; 26 < ML; 50 < ML) of ML by use 57 mammals which have real life-history traits data (**Table S15-17**).

### Gene set enrichment analysis

We use ‘Polysel’ (*46, 47*) for gene set enrichment analysis which is possible to detect pathways containing pleiotropic signals. In addition, other variables (such as gene length, number of species, and genetic distance) can also be used to adjust statistical variables. For the species-specific expression, we used the species-specific expression index (Tau) as the gene score (SUMSTAT) to detect species-specific expression pathways and ubiquitous expression (1 - Tau) pathways. For gene expression variation, we used the coefficients in the PGLS regression as SUMSTAT to enrich genes that are positively related to longevity (SUMSTAT of negatively related genes is set to 0) and negatively related genes (SUMSTAT of positively related genes is set to 0 and converted to Absolute value) to detect longevity-related pathways with genetic minor effects. Since gene expression is mostly related to gene length or species number, we used the function ‘RescaleBins’ to adjust SUMSTAT. We used ‘ks.test’ in R to check whether the gene score (SUMSTAT) is normally distributed or not. If not, a random data set was generated to construct an empirical distribution.

### Gene category collection

We collected different types of gene sets from various sources for comparison. The essential genes were constructed based on the probability of intolerance to loss of function, which is the pLI score (*150*). The score data comes from ExAC version 0.3.1 (https://gnomad.broadinstitute.org/). Genes with pLI> 0.9 are defined as essential genes. The list of genes associated with human inherited disease was obtained from the manually curated HGMD (PRO 17.1) (*151*). Aging genes were obtained from the GenAge database (*1*) and determined based on experimental evidence from humans and model organisms. They included genes related to the basic human aging process as well as genes related to lifespan. According to the homology relationship, the respective gene ID numbers were converted into human gene ID numbers. The Haploid Insufficiency (HI) score from previous studies (*152*) was used to quantify the degree of haploid deficiency in human genes. After sorting in descending order, we defined genes greater than the first quartile as haplo-insufficient genes, and genes less than the fourth quartile as haplo-sufficient genes. Finally, the phylogenetic age of mammalian genes was retrieved from the GenTree database (http://gentree.ioz.ac.cn/) (*153*). We divided genes into two groups based on genetic age: (i) those genes that appear after therian, mammalian, vertebrate, or quadrupedal ancestors (genes are defined as relatively young) and (ii) those that appear earlier than bone vertebrates Genes (defined as relatively old genes).

## Data and code availability

Raw RNA-seq data for liver and kidney of this study are available from Sciencedb with doi number 10.11922/sciencedb.01196. Raw RNA-seq data for brain are available from Sciencedb with doi number 10.11922/sciencedb.01197. The SRA ids of the data retrieved from NCBI are listed in **Table S1**. The code of this study is available from github (https://github.com/liu-wq/expressionML).

## Acknowledgements

We thank Alice Hughes, Xing Chen and Yanhua Chen for the experience and technology of sample collection. We thank Pengcheng Wang, Xiaoxiao Zhang, Zhan Zhang, and Qi Pan for supporting sample collection and preparation. This project was supported by National Natural Science Foundation of China (82050002) and the Beijing Natural Sciences Foundation (JQ19022). VNG is supported by grants from the National Institute on Aging.

## Author contributions

X.Z. conceived the study and designed the project. W.L. and P.Z. managed the project. W.L., P.Z., M.L., Z.L., Y.Y., G.L., X.J. and X.W. collected samples. W.L., P.Z., M.L., Z.L., Y.Y., J.D. and J.Y. prepared samples and performed RNA extraction. W.L. performed transcriptome assembly, annotation and bioinformatics analysis. W.L. and P.Z. performed life-history collection and imputation. W.L. and P.Z. discussed the data. W.L. wrote the manuscript with contributions from X.Z., M.L., V.N.G., I.S., L.W. and A.K.. All authors contributed to data interpretation.

## Competing interests

The authors declare no competing interests.

## Notes

### Competing Interest Statement

The authors have declared no competing interest.

