## Supplemental Tables and Figures for "Evolutionary transcriptomics reveals longevity mostly driven by polygenic and indirect selection in mammals": SIfile.pdf

**The PDF file includes:**

**Figs. S1 to S12**

**Legends for Tables S1 to S22**

### SI Figures

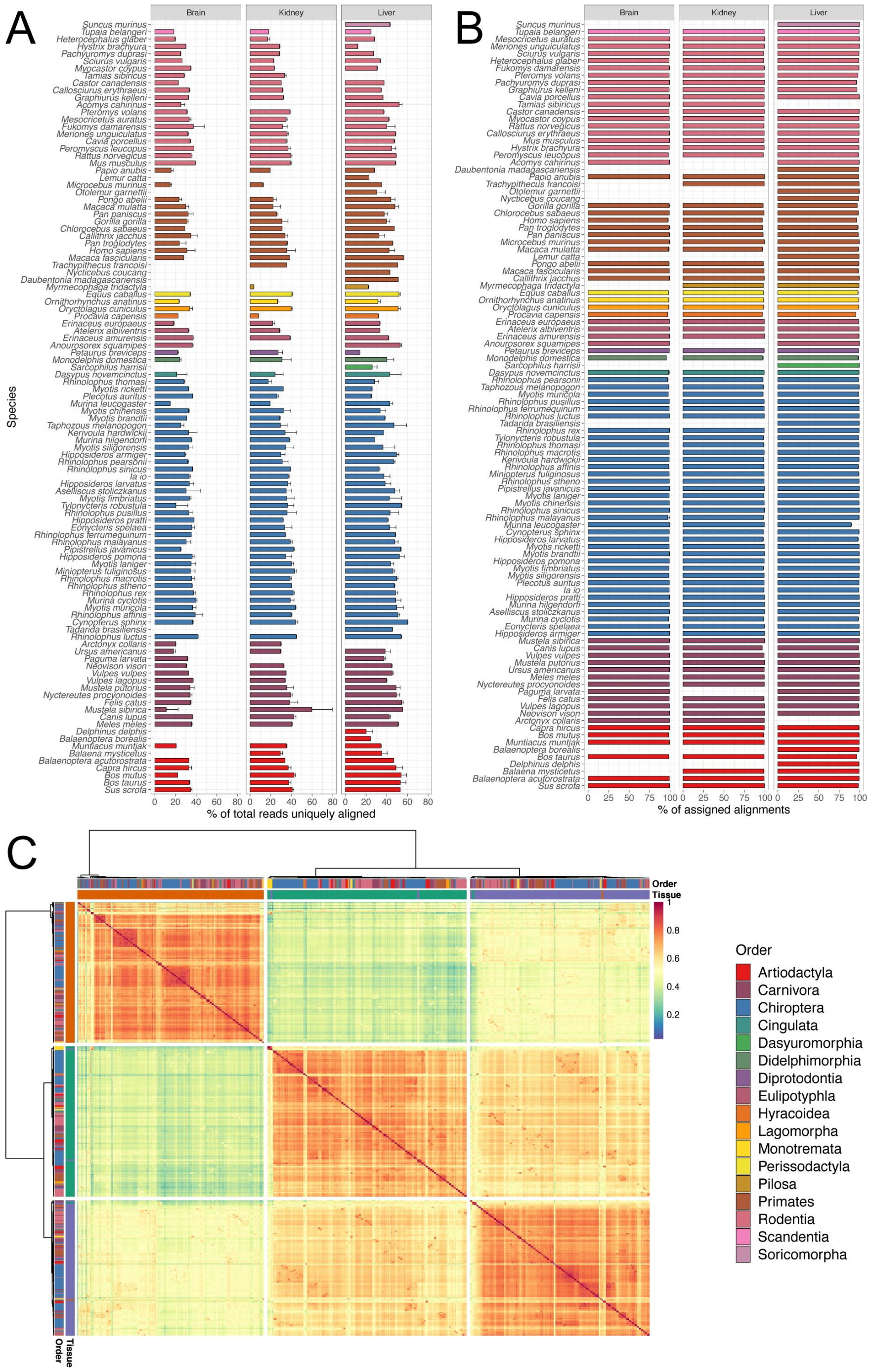


**Fig. S1 Sample alignment statistics and inter-sample correlation.** (*A*) RNA-seq unique mapping rates in each tissue. (*B*) The ratio of reads assigned to genes in each tissue. (*C*) Heatmap of correlation coefficient between samples. All group colors represent different order.

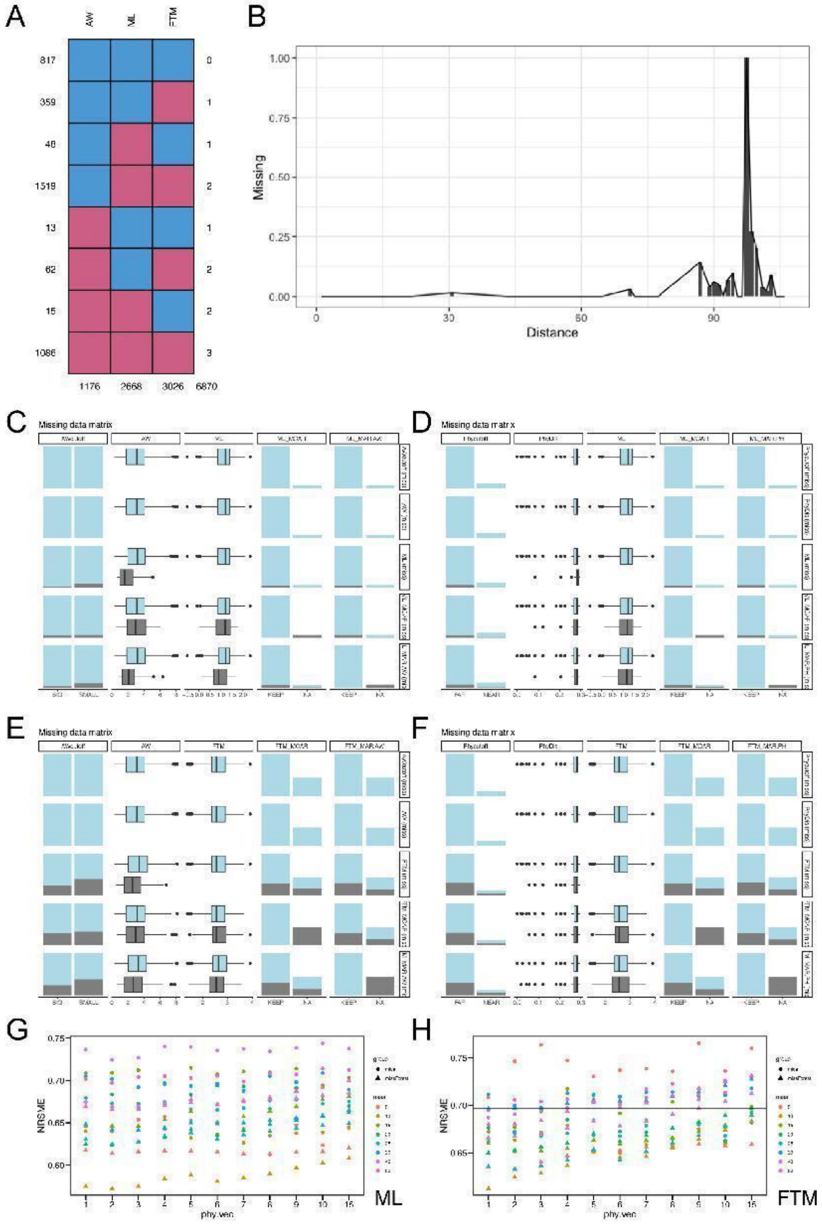


**Fig S2 Life history data imputation.** (*A*) Distribution of missing values in different life histories. (*B*) Distribution of missing values with increasing genetic distance from humans. (*C*) Correlation between missing value of maximum lifespan and adult weight. (*D*) Correlation between missing value of maximum life span and genetic distance from human. (*E*) Correlation between missing value of female time to maturity and adult weight. (*F*) Correlation between missing value of female time to maturity and genetic distance from human. (*G*) For the maximum lifespan, the gradient estimation accuracy comparison of the number of phylogenetic vectors added in the model. The colors represent different proportions of missing values, and the shapes represent two imputation methods. (*H*) For the female time to maturity, the gradient estimation accuracy comparison of the number of phylogenetic vectors added in the model. The colors represent different proportions of missing values, and the shapes represent two imputation methods.

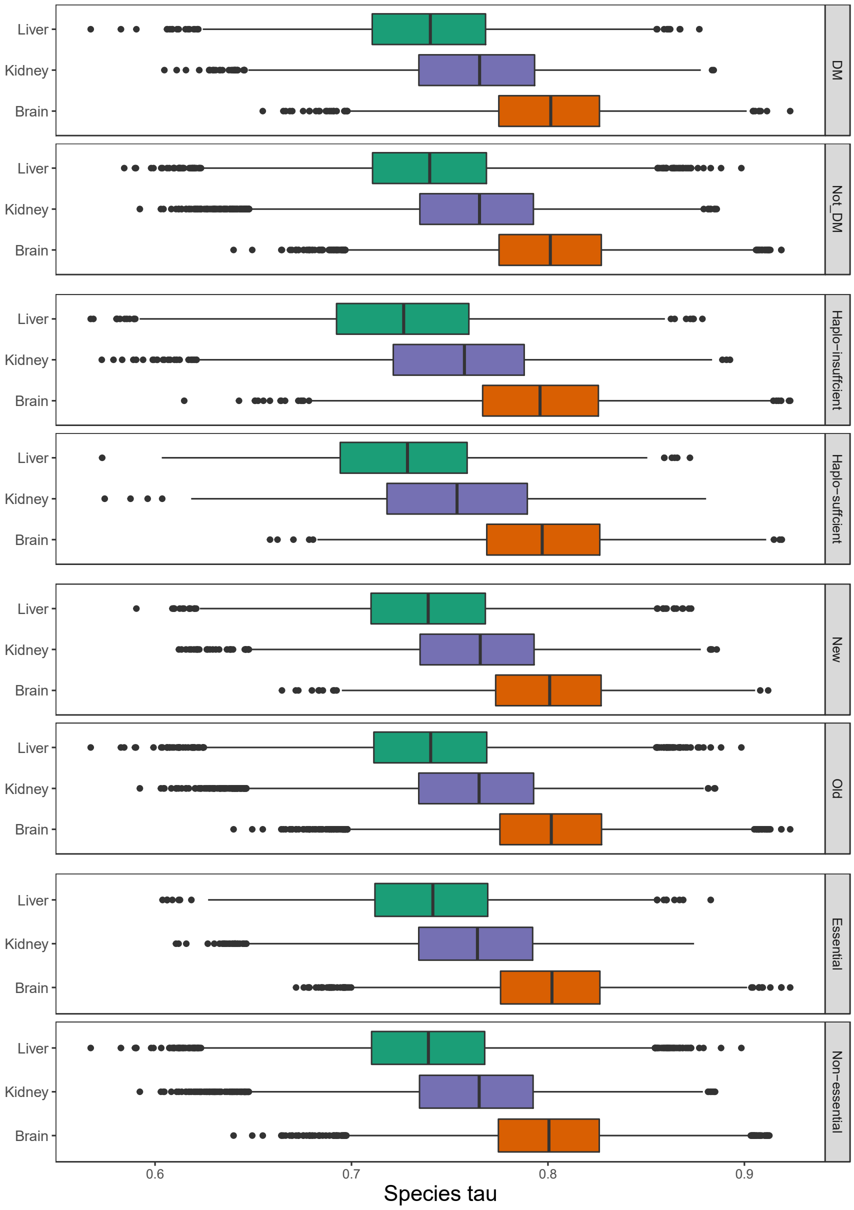


**Fig. S3 Distribution of the species-specific index Tau of gene expression in different gene category.**

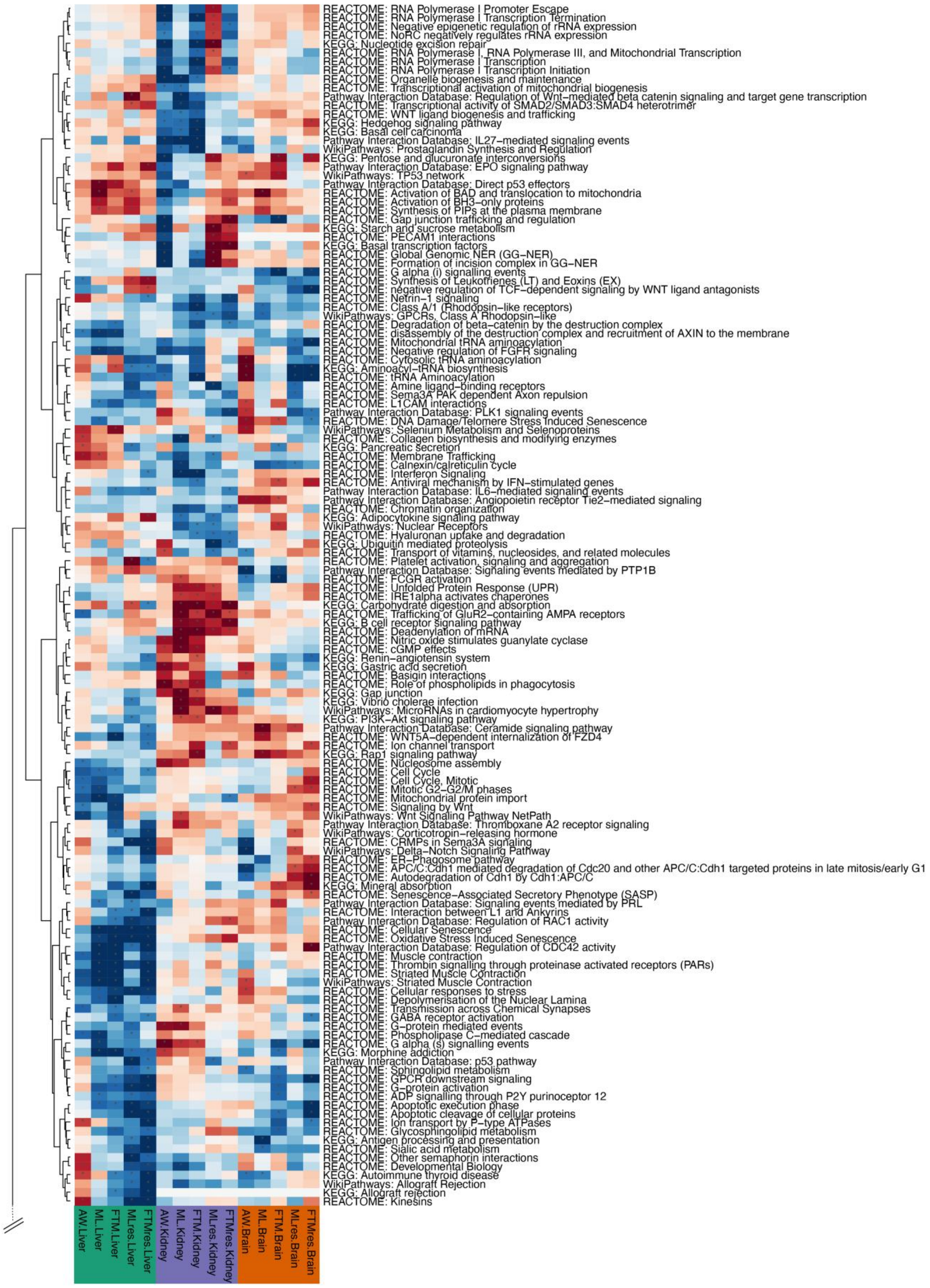
REACTOME: RNA Polymerase I Promoter Escape
REACTOME: RNA Polymerase I Transcription Termination
REACTOME: Negative epigenetic regulation of rRNA expression
REACTOME: NoRC negatively regulates rRNA expression
KEGG: Nucleotide excision repair
REACTOME: RNA Polymerase I, RNA Polymerase III, and Mitochondrial Transcription
REACTOME: RNA Polymerase I Transcription
REACTOME: RNA Polymerase I Transcription Initiation
REACTOME: Organelle biogenesis and maintenance
REACTOME: Transcriptional activation of mitochondrial biogenesis
Pathway Interaction Database: Regulation of Wnt−mediated beta catenin signaling and target gene transcription
REACTOME: Transcriptional activity of SMAD2/SMAD3:SMAD4 heterotrimer
REACTOME: WNT ligand biogenesis and trafficking
KEGG: Hedgehog signaling pathway
KEGG: Basal cell carcinoma
Pathway Interaction Database: IL27−mediated signaling events
WikiPathways: Prostaglandin Synthesis and Regulation
KEGG: Pentose and glucuronate interconversions
Pathway Interaction Database: EPO signaling pathway
WikiPathways: TP53 network
Pathway Interaction Database: Direct p53 effectors
REACTOME: Activation of BAD and translocation to mitochondria
REACTOME: Activation of BH3−only proteins
REACTOME: Synthesis of PIPs at the plasma membrane
REACTOME: Gap junction trafficking and regulation
KEGG: Starch and sucrose metabolism
REACTOME: PECAM1 interactions
KEGG: Basal transcription factors
REACTOME: Global Genomic NER (GG−NER)
REACTOME: Formation of incision complex in GG−NER
REACTOME: G alpha (i) signalling events
REACTOME: Synthesis of Leukotrienes (LT) and Eoxins (EX)
REACTOME: negative regulation of TCF−dependent signaling by WNT ligand antagonists
REACTOME: Netrin−1 signaling
REACTOME: Class A/1 (Rhodopsin−like receptors)
WikiPathways: GPCRs, Class A Rhodopsin−like
REACTOME: Degradation of beta−catenin by the destruction complex
REACTOME: disassembly of the destruction complex and recruitment of AXIN to the membrane
REACTOME: Mitochondrial tRNA aminoacylation
REACTOME: Negative regulation of FGFR signaling
REACTOME: Cytosolic tRNA aminoacylation
KEGG: Aminoacyl−tRNA biosynthesis
REACTOME: tRNA Aminoacylation
REACTOME: Amine ligand−binding receptors
REACTOME: Sema3A PAK dependent Axon repulsion
REACTOME: L1CAM interactions
Pathway Interaction Database: PLK1 signaling events
REACTOME: DNA Damage/Telomere Stress Induced Senescence
WikiPathways: Selenium Metabolism and Selenoproteins
REACTOME: Collagen biosynthesis and modifying enzymes
KEGG: Pancreatic secretion
REACTOME: Membrane Trafficking
REACTOME: Calnexin/calreticulin cycle
REACTOME: Interferon Signaling
REACTOME: Antiviral mechanism by IFN−stimulated genes
Pathway Interaction Database: IL6−mediated signaling events
Pathway Interaction Database: Angiopoietin receptor Tie2−mediated signaling
REACTOME: Chromatin organization
KEGG: Adipocytokine signaling pathway
WikiPathways: Nuclear Receptors
REACTOME: Hyaluronan uptake and degradation
KEGG: Ubiquitin mediated proteolysis
REACTOME: Transport of vitamins, nucleosides, and related molecules
REACTOME: Platelet activation, signaling and aggregation
Pathway Interaction Database: Signaling events mediated by PTP1B
REACTOME: FCGR activation
REACTOME: Unfolded Protein Response (UPR)
REACTOME: IRE1alpha activates chaperones
KEGG: Carbohydrate digestion and absorption
REACTOME: Trafficking of GluR2−containing AMPA receptors
KEGG: B cell receptor signaling pathway
REACTOME: Deadenylation of mRNA
REACTOME: Nitric oxide stimulates guanylate cyclase
REACTOME: cGMP effects
KEGG: Renin−angiotensin system
KEGG: Gastric acid secretion
REACTOME: Basigin interactions
REACTOME: Role of phospholipids in phagocytosis
KEGG: Gap junction
KEGG: Vibrio cholerae infection
WikiPathways: MicroRNAs in cardiomyocyte hypertrophy
KEGG: PI3K−Akt signaling pathway
Pathway Interaction Database: Ceramide signaling pathway
REACTOME: WNT5A−dependent internalization of FZD4
REACTOME: Ion channel transport
KEGG: Rap1 signaling pathway
REACTOME: Nucleosome assembly
REACTOME: Cell Cycle
REACTOME: Cell Cycle, Mitotic
REACTOME: Mitotic G2−G2/M phases
REACTOME: Mitochondrial protein import
REACTOME: Signaling by Wnt
WikiPathways: Wnt Signaling Pathway NetPath
Pathway Interaction Database: Thromboxane A2 receptor signaling
WikiPathways: Corticotropin−releasing hormone
REACTOME: CRMPs in Sema3A signaling
WikiPathways: Delta−Notch Signaling Pathway
REACTOME: ER−Phagosome pathway
REACTOME: APC/C:Cdh1 mediated degradation of Cdc20 and other APC/C:Cdh1 targeted proteins in late mitosis/early G1
REACTOME: Autodegradation of Cdh1 by Cdh1:APC/C
KEGG: Mineral absorption
REACTOME: Senescence−Associated Secretory Phenotype (SASP)
Pathway Interaction Database: Signaling events mediated by PRL
REACTOME: Interaction between L1 and Ankyrins
Pathway Interaction Database: Regulation of RAC1 activity
REACTOME: Cellular Senescence
REACTOME: Oxidative Stress Induced Senescence
Pathway Interaction Database: Regulation of CDC42 activity
REACTOME: Muscle contraction
REACTOME: Thrombin signalling through proteinase activated receptors (PARs)
REACTOME: Striated Muscle Contraction
WikiPathways: Striated Muscle Contraction
REACTOME: Cellular responses to stress
REACTOME: Depolymerisation of the Nuclear Lamina
REACTOME: Transmission across Chemical Synapses
REACTOME: GABA receptor activation
REACTOME: G−protein mediated events
REACTOME: Phospholipase C−mediated cascade
REACTOME: G alpha (s) signalling events
KEGG: Morphine addiction
Pathway Interaction Database: p53 pathway
REACTOME: Sphingolipid metabolism
REACTOME: GPCR downstream signaling
REACTOME: G−protein activation
REACTOME: ADP signalling through P2Y purinoceptor 12
REACTOME: Apoptotic execution phase
REACTOME: Apoptotic cleavage of cellular proteins
REACTOME: Ion transport by P−type ATPases
REACTOME: Glycosphingolipid metabolism
KEGG: Antigen processing and presentation
REACTOME: Sialic acid metabolism
REACTOME: Other semaphorin interactions
REACTOME: Developmental Biology
KEGG: Autoimmune thyroid disease
WikiPathways: Allograft Rejection
KEGG: Allograft rejection
REACTOME: Kinesins
AW.Liver
ML.Liver
FTM.Liver
MLres.Liver
FTMres.Liver
AW.Kidney
ML.Kidney
FTM.Kidney
MLres.Kidney
FTMres.Kidney
AW.Brain
ML.Brain
FTM.Brain
MLres.Brain
FTMres.Brain

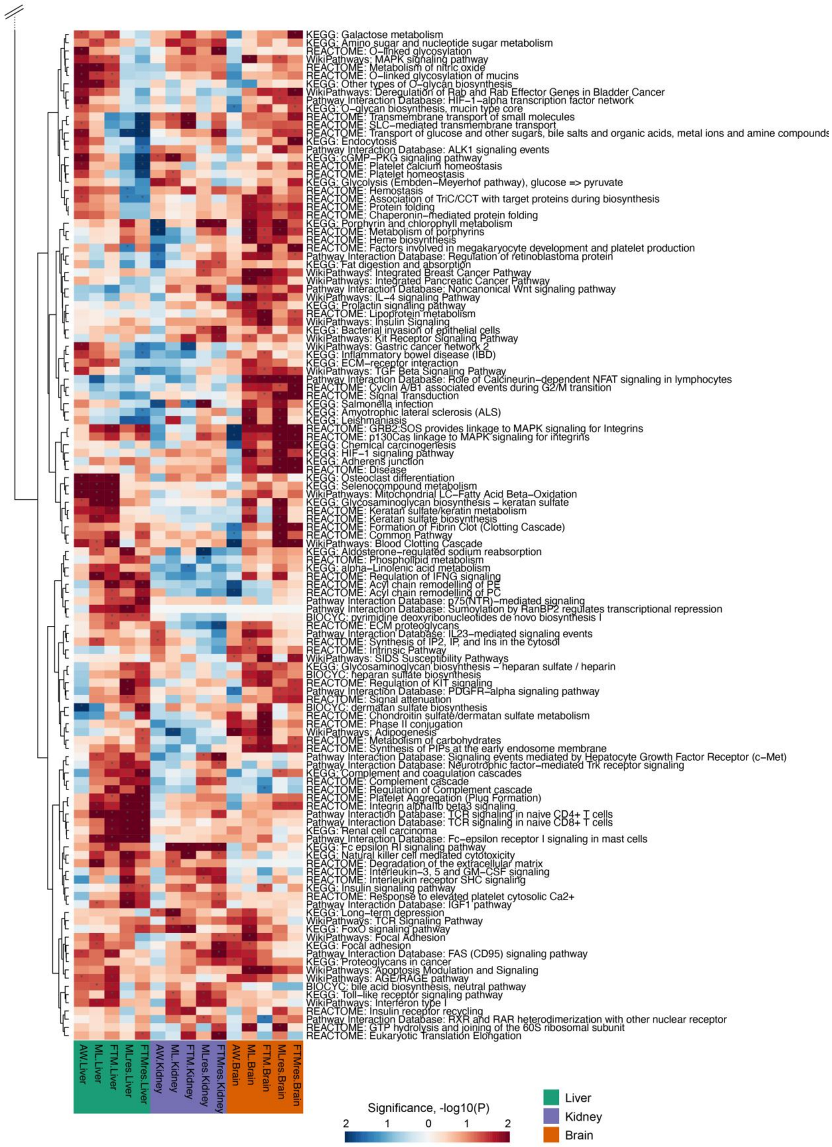


**Fig. S4 Gene set enrichment analysis heat map for all traits (prepruning).** Represents a clustering heat map of pathways enriched by longevity-related genes and AW-related genes. The color intensity indicates the degree of significance, and the *P* value has undergone a -log10 transformation. Each row represents a different pathway, and each column represents related longevity traits (marked at the bottom). Among them, red is positive correlation, and blue is negative correlation.

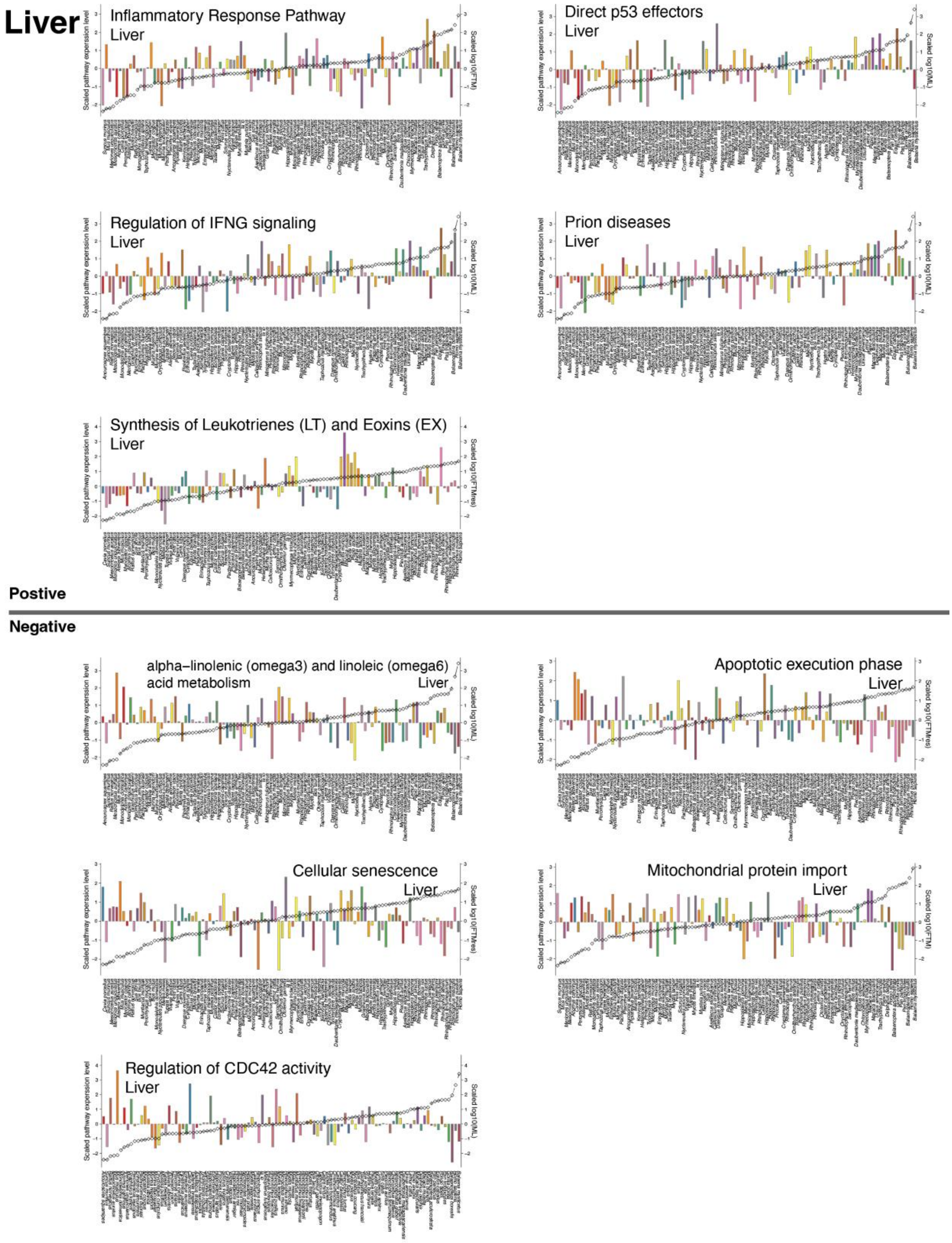


**Fig. S5 Top pathway for gene set enrichment in liver.** The y-axis on the left represents the sum of the expression levels of all genes in the alas pathway of each species. The y-axis on the right represents the value of longevity-related traits are centered at 0 on log10 scale. The x-axis is the species name, and the ranking increases with the longevity-related traits. The upper part of the gray line is the positive correlation pathway, and the lower part is the negative correlation pathway.

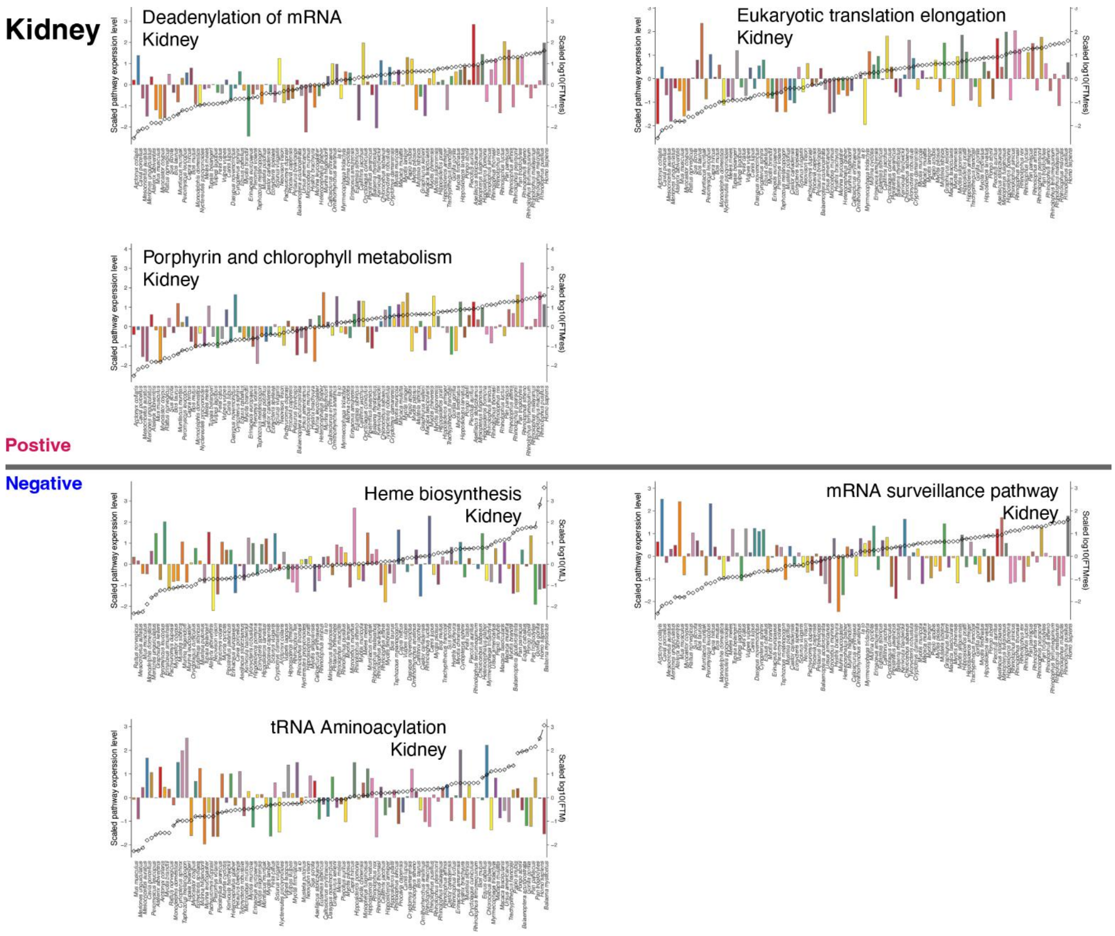


**Fig. S6 Top pathway for gene set enrichment in kidney.** The y-axis on the left represents the sum of the expression levels of all genes in the alas pathway of each species. The y-axis on the right represents the value of longevity-related traits are centered at 0 on log10 scale. The x-axis is the species name, and the ranking increases with the longevity-related traits. The upper part of the gray line is the positive correlation pathway, and the lower part is the negative correlation pathway.

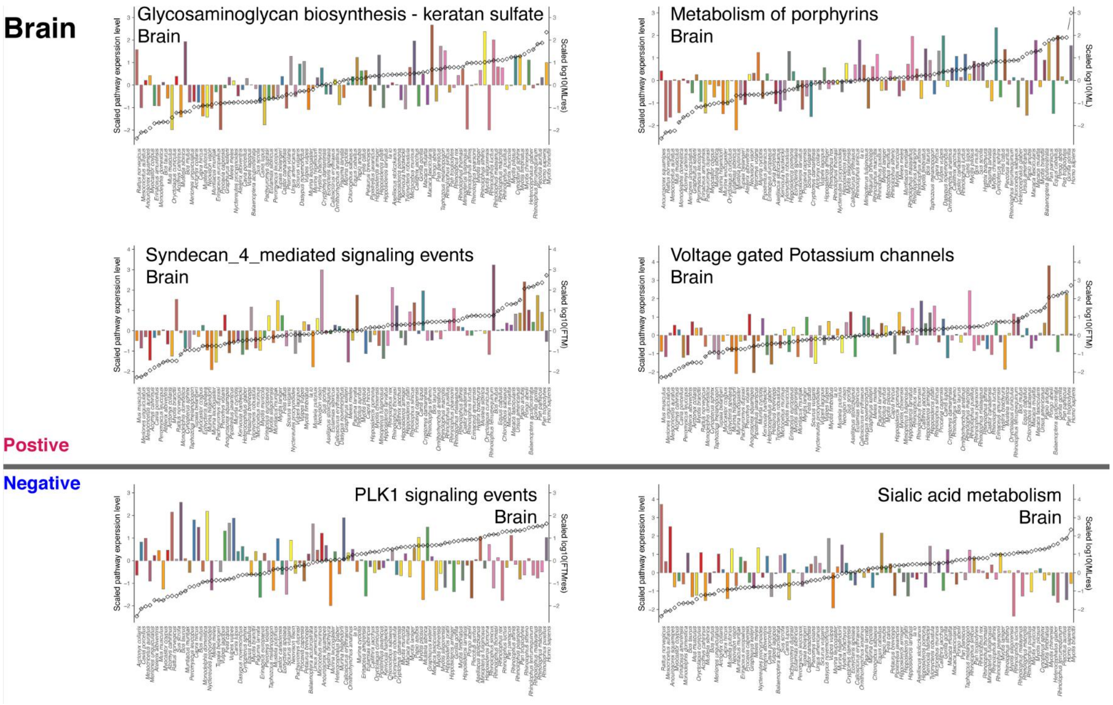


**Fig. S7 Top pathway for gene set enrichment in brain.** The y-axis on the left represents the sum of the expression levels of all genes in the alas pathway of each species. The y-axis on the right represents the value of longevity-related traits are centered at 0 on log10 scale. The x-axis is the species name, and the ranking increases with the longevity-related traits. The upper part of the gray line is the positive correlation pathway, and the lower part is the negative correlation pathway.

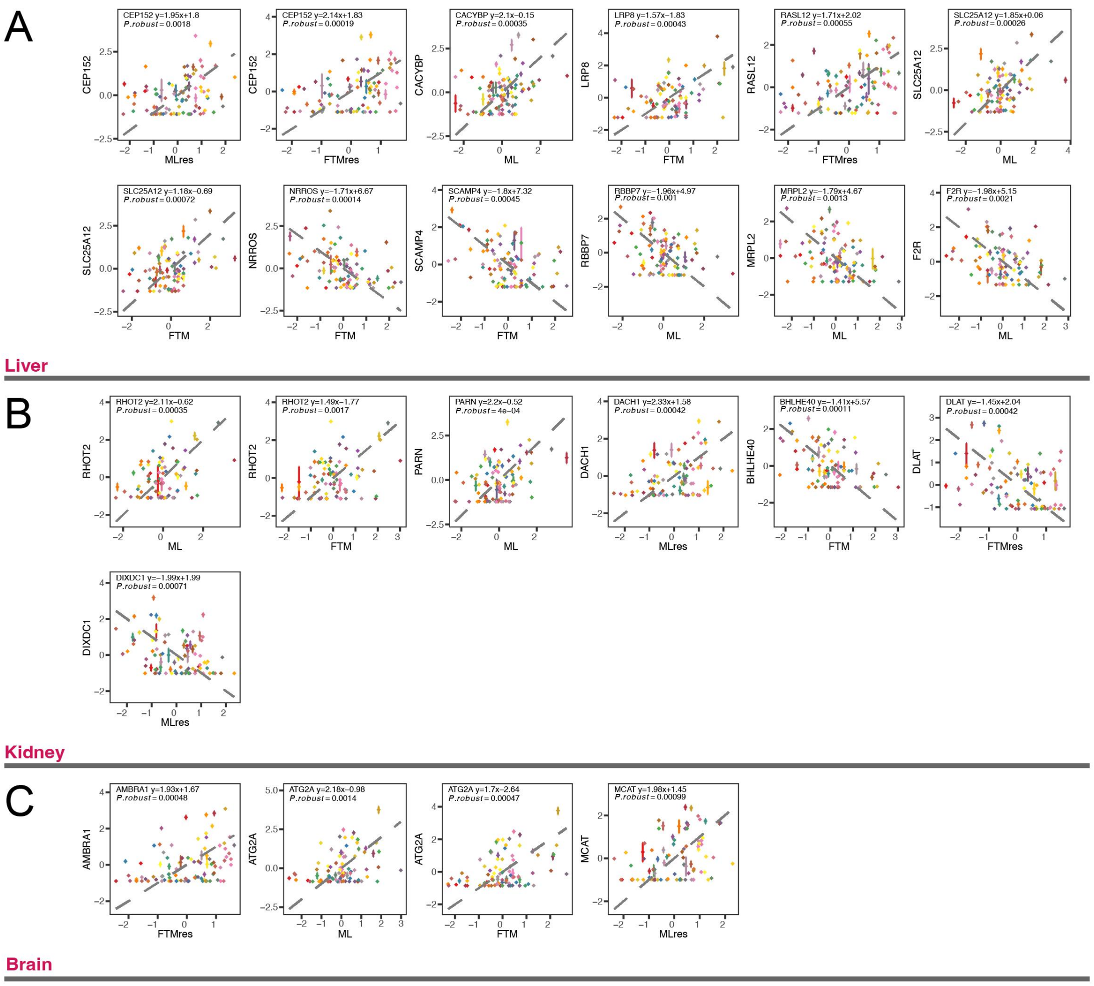


**Fig. S8 Selected genes with significant correlation to longevity.** (*A*) Longevity-related genes selected in the liver. (*B*) Longevity-related genes selected in the kidney. (*C*) Longevity-related genes selected in the brain. In each figure, the y-axis is the scaled expression level of each gene with 0 as the center, and the x-axis is the longevity traits (ML: maximum lifespan; FTM: female time to maturity; MLres and FTMres: ML and FTM residuals adjusted for adult weight, respectively). Potential outliers have been removed. The optimal phylogenetic regression equation, *P* value and $r^2$ are marked in the figure.

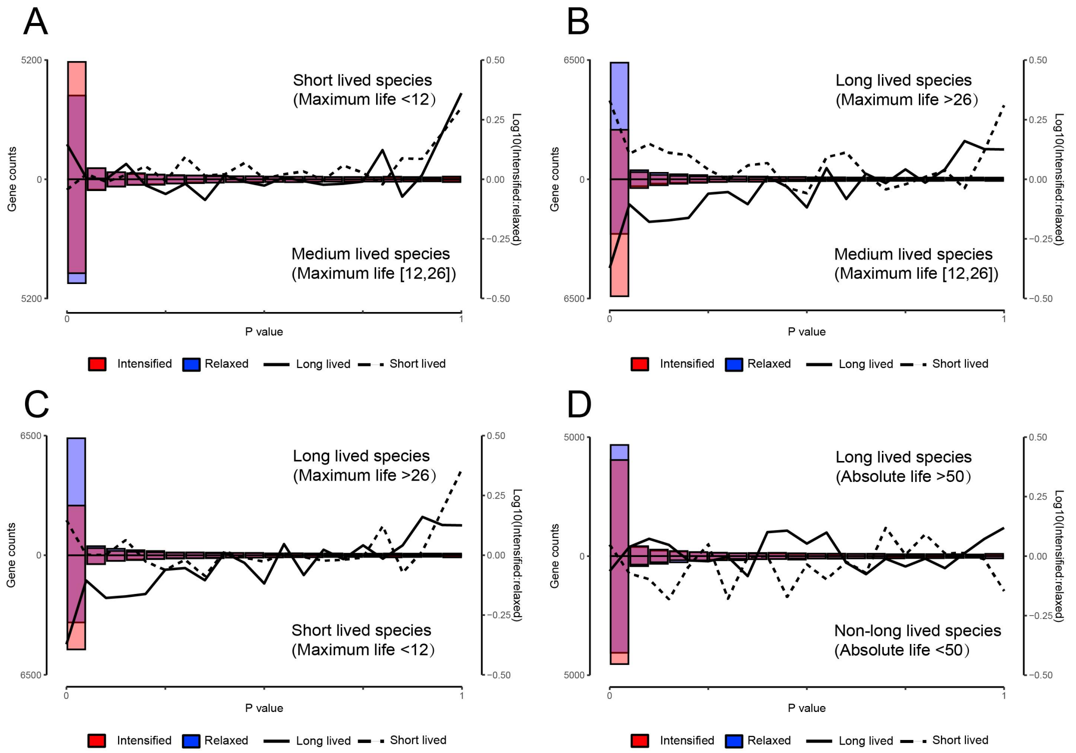


**Fig. S9 Relaxation of Selection in different lifespan interval.** P value distributions of the RELAX tests of long-lived species (ML > 26 or ML > 50), medium-lived species (12 <= ML <= 26), short-lived species (ML < 12) and non-long-lived species (ML < 50). (*A*) Contrast between short-lived species and medium-lived species, (*B*) Contrast between long-lived species and medium-lived species, (*C*) Contrast between long-lived species and short-lived species, and (*D*) Contrast between long-lived species and non-long-lived species are shown. Blue bars, genes under relaxed selection (k < 1); red bars, intensified genes (k > 1). Dotted (short-lived) and solid (long-lived) lines are the log ratios of k > 1: k < 1 gene counts.

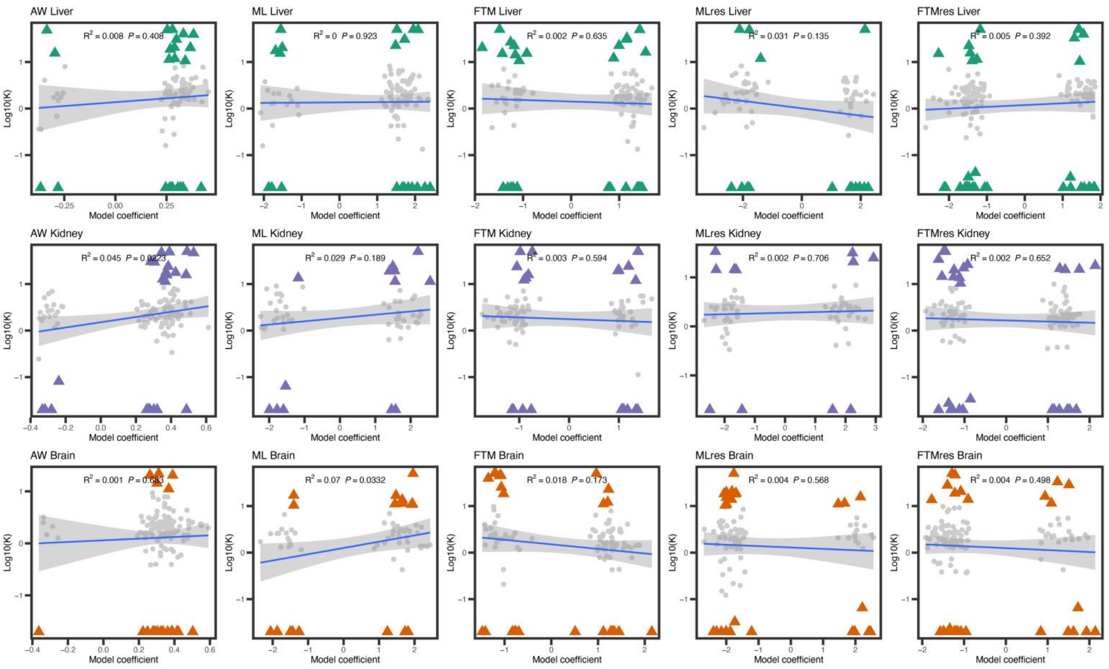


**Fig. S10 The relationship between the index of relaxed choice (K) and the regression coefficient.** In each figure, the regression coefficients of longevity-related genes for each longevity trait in different tissues and the index of relaxed selection (K) are shown. The x-axis represents the regression coefficient, and the y-axis represents the relaxation selection index (K) on log10 scale. The points in the figure are longevity-related genes and the selection pressure has also changed significantly. Among them, gray represents longevity-related genes with low selection intensity, and red are longevity-related genes with high selection intensity. The regression line, 95% confidence interval, *P* value and $r^2$ are marked in the figure.

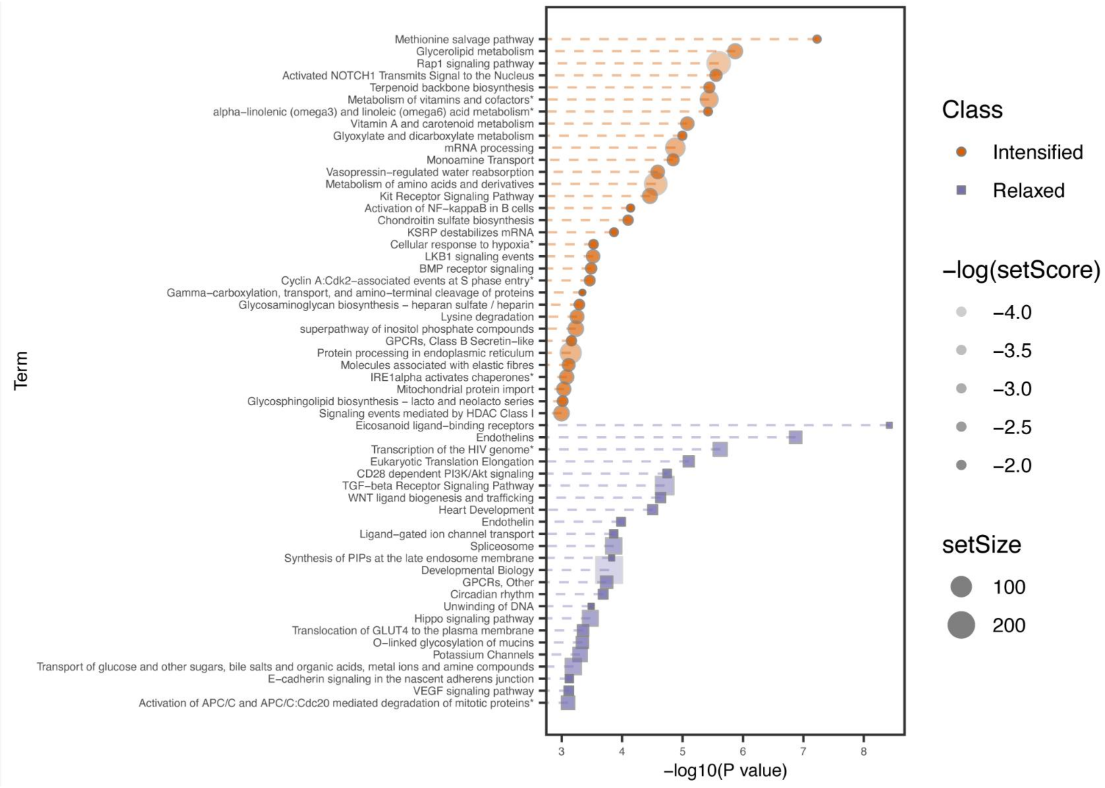


**Fig. S11 Gene set enrichment of genes under relaxed or strengthened selection.** Polysel gene set enrichment analysis of genes under relaxed or strengthened selection. The x-axis is the *P* value on -log10 scale. Different selection uses different shapes and colors. The shade of the color represents the SUMSTAT, and the size of the dot represents the size of the gene set.

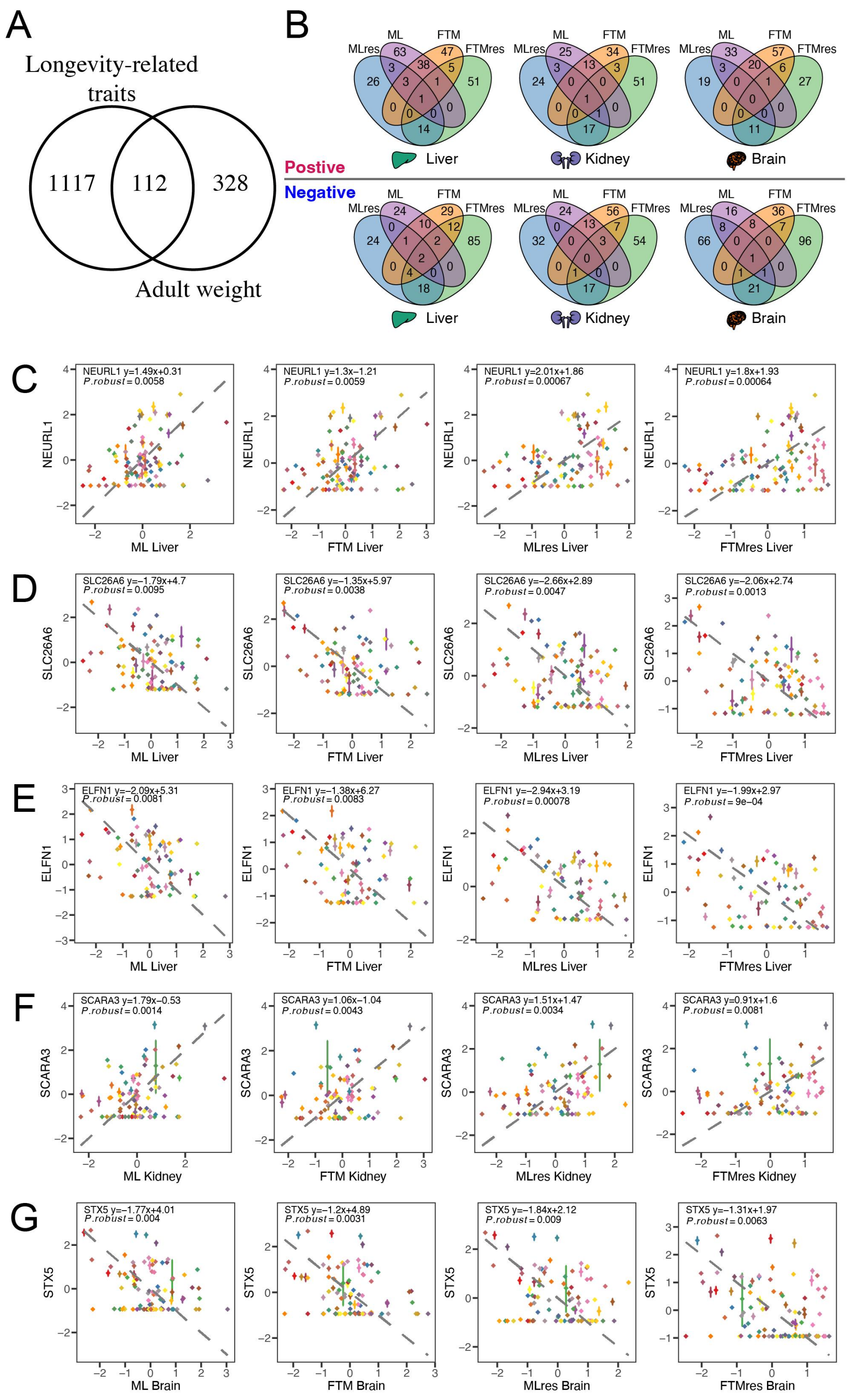


**Fig. S12 Overlap of longevity-related genes.** (*A*) Venn diagram of overlap between longevity-related genes and adult weight-related genes. Genes that are significant in the four longevity traits (*B*) Venn diagrams of each tissue for positive/negative longevity-related genes. (*C*) *NEURL* show positive correlation with the longevity traits in the liver. (*D*)*SLC26A6* and (*E*) *ELFN1* show negative correlation with the longevity traits in the liver. (*F*) *SCARA3* show positive correlation with the longevity traits in the kidney. (*G*) STX5 show negative correlation with the longevity traits in the brain. In each figure, the y-axis is the scaled expression level of each gene with 0 as the center, and the x-axis is the longevity traits (ML: maximum lifespan; FTM: female time to maturity; MLres and FTMres: ML and FTM residuals adjusted for adult weight, respectively). Potential outliers have been removed. The optimal phylogenetic regression equation, $P_{robust}$ is marked in the figure.

### Supplementary Tables

**Table S1. The sample information and sequencing platform of 106 mammals analyzed in the present study.**

**Table S2. The classification and life histories of 106 mammals analyzed in the present study.**

[1]Life histories that were imputed and estimated in this study.

**Table S3. The species-specific expression index (tau) for each gene in three organs.**

[1] The number of species used in estimation of Tau.

[2] The species with the highest expression of the gene.

[3] The top 10 species with the highest expression of this gene

**Table S4. The significant scoring gene sets post-pruning for species-specific expression index.**

[1] Number of genes in pathway.

[2] SUMSTAT score for each pathway.

**Table S5. The significant scoring gene sets pre-pruning for species-specific expression index.**

[1] Number of genes in pathway.

[2] SUMSTAT score for each pathway.

**Table S6. Phylogenetic regression of gene expression against longevity traits.**

[1] P value of phylogenetic regression optimal model

[2] The P value adjusted by the two-step method in PGLS, including *P.robust* and *P.max*.

[3] Number of species used in PGLS.

**Table S7. The significant scoring gene sets post-pruning for the coefficient of each longevity traits.**

[1] Number of genes in pathway.

[2] SUMSTAT score for each pathway.

**Table S8. The significant scoring gene sets pre-pruning for the coefficient of each longevity traits.**

[1] Number of genes in pathway.

[2] SUMSTAT score for each pathway.

**Table S9. Phylogenetic regression of gene expression against longevity traits in Primates, Rodents, and Chiroptera.**

**Table S10. The significant scoring gene sets post-pruning for the coefficient of each longevity traits in Primates, Rodents, and Chiroptera**

[1] Number of genes in pathway.

[2] SUMSTAT score for each pathway.

**Table S11. The significant scoring gene sets pre-pruning for the coefficient of each longevity traits in Primates, Rodents, and Chiroptera.**

[1] Number of genes in pathway.

[2] SUMSTAT score for each pathway.

**Table S12. Functional annotation of gene mutation tolerance and adaptability.**

**Table S13. Genes under relax and intensified selection in long-lived animals estimated with RELAX.**

[1] Relaxation parameters.

[2] Three ω parameters and the relative proportion of sites in reference branch.

[3] Three ω parameters and the relative proportion of sites in foreground branch.

**Table S14. The statistic of number of genes identified in RELAX and with expression associated with longevity traits.**

[1] Number of genes positively or negatively related to lifespan under intensified selection and the number of genes with strong selection (in brackets).

[2] Number of genes positively or negatively related to lifespan under relaxed selection and the number of genes with strong selection (in brackets).

**Table S15. Genes under relax and intensified selection in long-lived animals and medium-lived animals estimated with RELAX.**

[1] Relaxation parameters.

[2] Three ω parameters and the relative proportion of sites in reference branch.

[3] Three ω parameters and the relative proportion of sites in foreground branch.

**Table S16. Genes under relax and intensified selection in medium-lived animals and short-lived animals estimated with RELAX.**

[1] Relaxation parameters.

[2] Three ω parameters and the relative proportion of sites in reference branch.

[3] Three ω parameters and the relative proportion of sites in foreground branch.

**Table S17. Genes under relax and intensified selection in long-lived animals and non-long-lived animals estimated with RELAX.**

[1] Relaxation parameters.

[2] Three ω parameters and the relative proportion of sites in reference branch.

[3] Three ω parameters and the relative proportion of sites in foreground branch.

**Table S18. The significant scoring gene sets post-pruning for the relaxation parameter (*k*) in RELAX.**

[1] Number of genes in pathway.

[2] SUMSTAT score for each pathway.

**Table S19. The significant scoring gene sets pre-pruning for the relaxation parameter (*k*) in RELAX.**

[1] Number of genes in pathway.

[2] SUMSTAT score for each pathway.

**Table S20. The statistics of alignments of RNA-seq reads.**

[1] Sequencing library size after filtering.

[2] Number of reads uniquely mapped to the orthologous gene set.

[3] Ratio of reads uniquely mapped to the orthologous gene set.

[4] Ratio of uniquely aligned reads assigned to genes.

**Table S21. Species classification in relaxed selection analysis.**

**Table S22. MCMCglmm summary.**
